# Pathogenesis of mtDNA point mutation m.10191T>C affecting complex I function is a multifactorial process leading to metabolic remodeling of mitochondria

**DOI:** 10.1101/2024.12.01.626247

**Authors:** Zeinab Alsadat Ahmadi, Alfredo Cabrera-Orefice, Marta Zaninello, Esther Barth, Richard Rodenburg, Elena I. Rugarli, Susanne Brodesser, Susanne Arnold, Ulrich Brandt

## Abstract

Inherited mitochondrial disorders are of multiple genetic origins and may lead to a broad range of frequently severe disease phenotypes. Yet, the correlation between molecular causes and clinical presentations is poorly understood. To address this conundrum, we thoroughly investigated the consequences of the well-known pathogenic mitochondrial DNA mutation m.10191T>C. The mutation changes serine-45 in subunit ND3 of respiratory chain complex I to proline and causes Leigh syndrome, which is one of the most devastating mitochondrial diseases. Human mitochondria carrying the mutation ND3^S45P^ retained 30-40% of complex I activity and oxidative phosphorylation capacity. In stark contrast, intact mutant cells exhibited only minimal oxygen consumption and a massively increased NADH/NAD^+^ ratio. Since the energy barrier for the Active/Deactive transition of complex I was reduced by ∼20 kJ·mol^-1^ in mutant cells, we concluded that complex I was shut-off by malfunctioning of an as yet unknown regulatory pathway. Comprehensive analysis of the mitochondrial complexome of cybrids, patient fibroblasts and muscle biopsies rendered other causes for the accumulation of NADH unlikely. The complexome datasets provide a rich resource for further studies to discover possible additional factors involved in regulating complex I. We propose that the derailed regulation of complex I is the main culprit leading to NADH accumulation and eventually the severity of the disease phenotype caused by mutation ND3^S45P^.

## Introduction

Mitochondrial dysfunction is an important factor in a wide range of pathologies including neurodegenerative disorders, diabetes, heart failure and cancer (Burke, 2017; Filosto *et al*, 2011; Morio *et al*, 2019; Rosca *et al*, 2013). Moreover, declining mitochondrial performance in general has been linked to biological aging (Jang *et al*, 2018). However, even in Parkinson’s disease, where a lot is known about the molecular processes involving mitochondria, it remains obscure why dopaminergic neurons are affected specifically, while other neurons, cell types and tissues remain largely unaffected (Borsche *et al*, 2021). Likewise, for inherited mitochondrial disorders, the correlation between molecular causes and affected organs or clinical presentations is poorly understood. It remains largely unknown, how mutations affecting the function of mitochondrial proteins lead to such a wide range of patient phenotypes from exercise intolerance to selective loss of vision or hearing, severe muscular or cardiac dysfunction to fatal neurological deficiencies (Fernandez-Vizarra & Zeviani, 2021; Ng *et al*, 2021).

Evidently, the multifaceted presentation of pathologies linked to mitochondria originates from their central roles in energy and intermediary metabolism, cofactor and lipid biogenesis, quality control, calcium homeostasis and intracellular signaling. While our knowledge about these molecular pathways has increased enormously in recent years, the chain of events leading from a given molecular cause to a specific pathological phenotype has remained enigmatic. This holds true even for the many inherited disorders caused by defects in components of the oxidative phosphorylation (OXPHOS) system, the most fundamental and best-studied mitochondrial process. Despite the intricate connections between energy metabolism and virtually all other mitochondrial functionalities, the prevailing paradigm still is that the predominant culprits leading to disease caused by OXPHOS defects are lowered ATP synthesis, high NADH/NAD^+^ ratio and increased production of deleterious reactive oxygen species (ROS). However, our recent finding illustrates that the complete elimination of OXPHOS by ablation of mitochondrial DNA (mtDNA) leads to widespread remodeling of the mitochondrial complexome and proteome (Guerrero-Castillo *et al*, 2021). Therefore, a broader, in-depth approach seems necessary to fully understand the pathogenic mechanisms unleashed by mitochondrial deficiencies.

Accordingly, we decided to track down the consequences of a well-known pathogenic mtDNA point mutation from the functional changes of the affected enzyme to remodeling of mitochondria as a whole. We chose m.10191T>C (ND3^S45P^), a prevalent mutation affecting the mitochondrially encoded subunit ND3 of respiratory complex I (Taylor *et al*, 2001).

Mitochondrial complex I (proton-pumping NADH:ubiquinone oxidoreductase) is the main entry point of electrons into the respiratory chain. It couples electron transfer from NADH to ubiquinone to the translocation of four protons across the inner mitochondrial membrane to drive ATP synthesis (Brandt, 2006; Zickermann *et al*, 2015). Besides its regular energy converting function, complex I is a major source of deleterious ROS (Dröse *et al*, 2016). Mammalian complex I, the largest component of the OXPHOS system, is of dual genetic origin. For its coordinated assembly (Guerrero-Castillo *et al*, 2017), seven mtDNA-encoded, hydrophobic, membrane-integral subunits need to be combined with an additional 37 subunits that are encoded by nuclear genes (Hirst, 2013) and, therefore, need to be imported into mitochondria after their synthesis by cytosolic ribosomes.

Mitochondrial dysfunction caused by mutations in mtDNA-encoded subunits of complex I prove to be exceptionally detrimental and result in a variety of severe clinical phenotypes. One of these mutations is the single base exchange m.10191T>C that replaces serine-45 of subunit ND3 with a proline. This renders the hydrophilic loop connecting the first and second α-helix of the subunit less flexible. The loop is a critical part of the ubiquinone reducing active site (Cabrera-Orefice *et al*, 2018; Zickermann *et al*, 2015) and a residue at its tip, cysteine-39, is a biochemical marker for the Active/Deactive (A/D) transition (Galkin *et al*, 2008), a still poorly understood regulatory mechanism of complex I (Grivennikova *et al*, 2020).

When occurring at a high mtDNA mutation load due to elevated heteroplasmy levels, mutation m.10191T>C (Taylor *et al*, 2001) causes Leigh syndrome (OMIM #256000), the most frequently observed manifestation of pediatric mitochondrial disease (Chang *et al*, 2020). Presenting progressive and severe neurological symptoms, most patients suffer from an early onset of disease and die at very young age (McFarland *et al*, 2004). Besides neurological phenotypes, such as mental retardation, developmental delay, ataxia and dystonia or spasticity, multisystemic defects in other organs, like heart, kidney and liver, were observed (Ruhoy & Saneto, 2014).

In addition to mutation m.10191T>C, several other mutations in mitochondrial as well as nuclear genes encoding subunits of complex I have been reported to cause Leigh syndrome (Fernandez-Vizarra & Zeviani, 2021). Genetic models of Leigh syndrome were generated by knocking out the nuclear encoded subunit NDUFS4 of complex I in mouse (Quintana *et al*, 2010) and *Yarrowia lipolytica* (Kahlhöfer *et al*, 2017). These models were studied extensively (Breuer *et al*, 2013; de Haas *et al*, 2017; Quintana & Hoth, 2012). Structural destabilization of the peripheral arm of complex I seems to lead to an increased production of ROS that are generally considered to be a critical factor for the progression of various mitochondrial disorders (Adjobo-Hermans *et al*, 2020; Kahlhöfer *et al*, 2017). However, while the complete removal of NDUFS4 from complex I results in a phenotype similar to Leigh syndrome in mice, it does not exactly reproduce any of the pathogenic mutations in humans.

Hence, we studied a specific point mutation of a complex I subunit gene that is responsible for a devastating and early fatal form of Leigh syndrome and that directly affects the catalytic function of complex I in human cells reasoning that this could allow a mechanistic interpretation of the molecular cause. Patients carrying a high load of mutation m.10191T>C are affected by very severe and rapidly progressing Leigh syndrome, but still exhibit significant levels of remaining complex I function. This suggested that factors other than complex I deficiency alone are relevant for the pathogenic process. Moreover, due to the importance of complex I in mitochondrial ROS production (Dröse *et al*, 2016) and the involvement of the loop of subunit ND3 in the A/D transition (Grivennikova *et al*, 2020), we expected to gain deeper insight into the pathogenic mechanism of this point mutation responsible for a devastating and early fatal form of Leigh syndrome.

## Results

### Mutation ND3^S45P^ hardly affects complex I abundance but impairs catalytic activity and inhibitor binding

We first analyzed the immediate impact of mutation ND3^S45P^ on complex I stability and function in human cybrids derived from 143B ρ0 cells that were homoplasmic for mutation m.10191T>C in comparison to cybrids containing 100% wildtype mtDNA as control (McFarland *et al*, 2004).

Complexome profiling analysis of mitochondria solubilized by digitonin showed hardly any effect on the abundance and stability of respirasomes that harbor most of complex I in human mitochondria (**Figure 1A**). Also, the pattern of assembly intermediates migrating at lower molecular masses, typically observed in mitochondria from proliferating cells (Guerrero-Castillo *et al*, 2017), was essentially unchanged. Assessment of the abundance of all five OXPHOS complexes based on label-free mass spectrometric quantification of their corresponding subunits (**Table S1**) revealed a slight reduction in the amounts of complex I, while complexes II to V were not changed significantly (**Figure 1B**). We concluded that mutation ND3^S45P^ had only minor, if any, impact on the assembly and stability of complex I.

**Figure 1:**
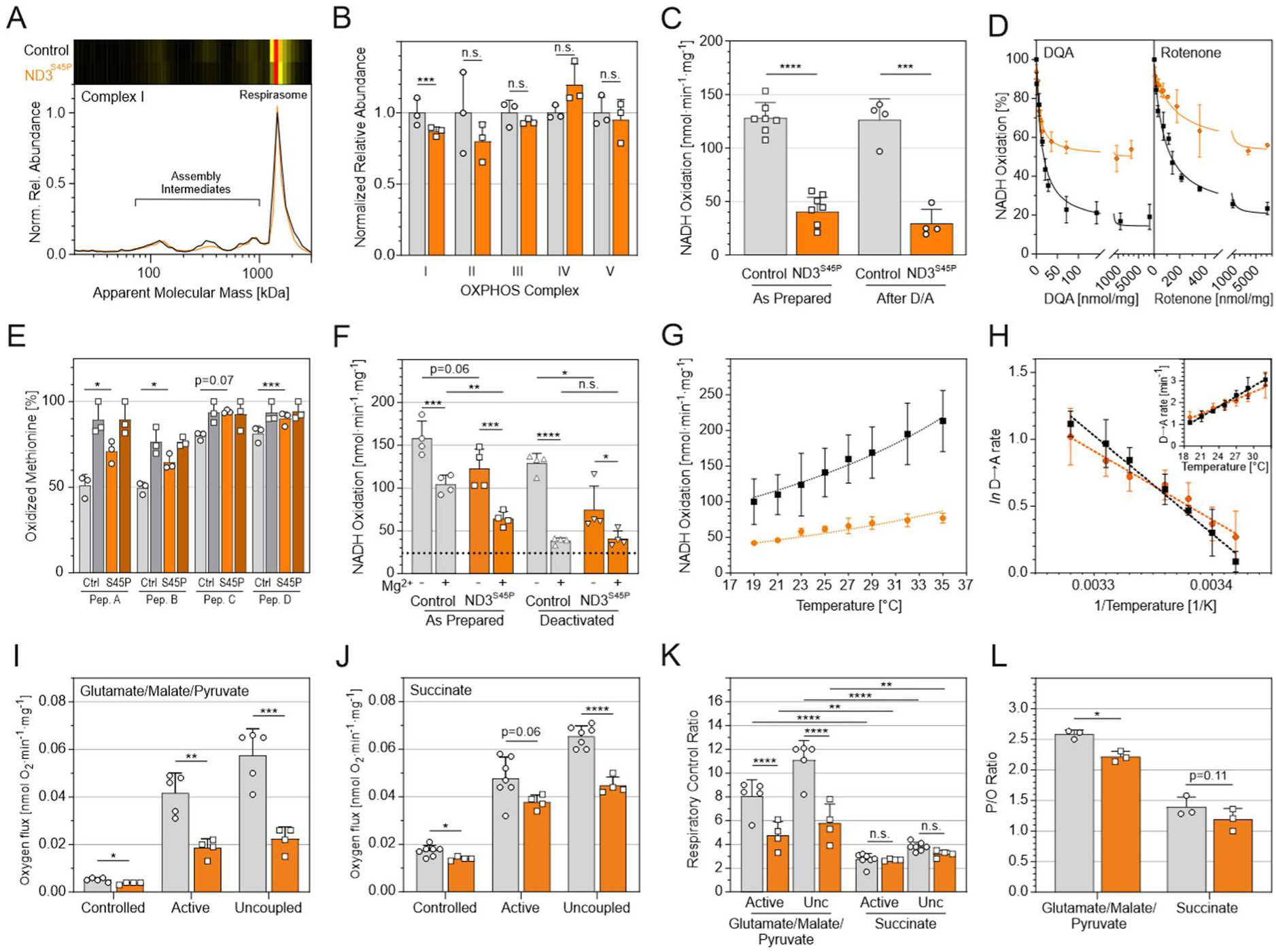
Biochemical analysis of complex I. Independently prepared batches of mitochondria from control (black/grey) and ND3^S45P^ (orange/brown) cybrids were analyzed. **A**, complexome migration profiles (bottom) and heatmaps (top) representing the average of all reliably detected subunits of complex I from three preparations of cybrids grown on glucose. **B**, relative abundance of OXPHOS complexes I-V as determined from the average of the abundances of all their consistently detected subunits. **C**, DQA-sensitive NADH oxidase activities in mitochondria as prepared (left) and after one cycle of deactivation followed by activation D/A (right). **D**, relative rates of NADH oxidation titrated with increasing amounts of the specific complex I inhibitors DQA (left) and rotenone (right) fitted by nonlinear regression. **E**, degree of methionine oxidation in peptides M^77^CLVEIEK (Pep. A), KTESIDVM^254^DAVGSNIVVSTR (Pep. B), KPM^473^VVLGSSALQR (Pep. C), ALSEIAGM^632^TLPY DTLDQVR (Pep. D) of complex I subunit NDUFS1 as determined from complexome profiling datasets of control and ND3^S45P^ cybrids on galactose (ϒ) and glucose (♦). **F**, total NADH oxidase activities in mitochondria as prepared (left) and after one cycle of deactivation followed by activation during the experiment (right) and in the presence or absence of 5 mM MgCl_2_. The dotted horizontal line indicates the average unspecific NADH activity not sensitive to complex I inhibitors in control cells. **G**, temperature dependence of NADH oxidase activity. **H**, Arrhenius plot of the temperature dependent rate of reactivation as deduced from the length of the lag phase of NADH oxidation (Insert). Mitochondrial respiration rates normalized to citrate synthase: Oxygen consumption rates with complex I linked substrates glutamate/malate/ pyruvate (**I**) and complex II linked substrate succinate (**J**). **K**, respiratory control ratios indicating the fold increase in respiration upon addition of ADP (active) or the uncoupler CCCP (Unc) to mitochondria in controlled state. **L**, P/O ratios as calculated from the amount of oxygen consumed to phosphorylate a defined amount of ADP. Data are presented as mean ± SD (n = 3-7) \**p*<0.05, \*\**p*<0.01, \*\*\**p*<0.001, \*\*\*\**p*<0.0001, n.s. not significant; two-way ANOVA over subunits (**B**), Student’s *t*-test (**C-F; I-L**) or fitted by nonlinear (**G**) or linear regression (**H**).

Next, we measured the effect of mutant ND3^S45P^ on complex I activity. The amino acid exchange lowered NADH oxidase activity sensitive to the complex I inhibitor 2-decyl-4-quinazolinylamine (DQA) in mitochondrial membranes by about 70%, and this was not changed significantly by subjecting the mitochondrial membranes to a cycle of deactivation followed by activation (Figure 1C). The amount of the complex I inhibitors DQA and rotenone needed to reduce the sensitive fraction of NADH oxidase activity by 50% (IC_50_) was only marginally affected: for DQA, IC_50_ decreased from 13 ± 2 to 7 ± 1 nmol·mg^-1^ (n = 3; *p*<0.01) and for rotenone it increased from 84 ± 11 to 160 ± 37 nmol·mg^-1^ (n = 3; *p*<0.05; Figure 1D). The inhibitor insensitive portion of total NADH oxidase activity was ∼40 nmol·min^-1^·mg^-1^ for DQA and ∼45 nmol·min^-1^·mg ^-1^ for rotenone. For both inhibitors, this corresponds to ∼20% and ∼55% of the uninhibited rate in control and mutant mitochondria, respectively.

### Mutation ND3^S45P^ moderately increases oxidation of NDUFS1 in cybrids on galactose

Increased production of ROS by complex I is assumed to frequently contribute to the pathogenesis linked to mitochondria (Dröse *et al*, 2016). Therefore, we explored whether mutation ND3^S45P^ led to an increased oxidative load in mitochondria. Due to their high reactivity and short lifetime, ROS predominantly have local impact and the actual modifications depend on the site where they are generated (Bleier *et al*, 2015). To identify changes specific to complex I dependent ROS production, we analyzed the complexome profiling datasets for oxidative modifications of amino acids in complex I subunits surrounding the FMN binding site, the major site of superoxide formation (Galkin & Brandt, 2005).

We identified four methionines in subunit NDUFS1 (M77, M254, M473, M632) that were 10-20% more oxidized in cybrids carrying mutation ND3^S45P^ than in controls suggesting a moderate increase in ROS production by complex I (Figure 1E). Interestingly, this increase was only observed when cells were forced to use OXPHOS by growing them on galactose instead of glucose media. On glucose, the levels of methionine oxidation were not at all affected by the mutation and remained consistently higher than those on galactose, reaching or exceeding the levels observed for mutant cells on galactose (Figure 1E). Moreover, while on galactose, the only detectable modification of methionine was sulfoxidation, on glucose ∼5% of M473 were further oxidized to methionine sulfone and ∼1% further to homocysteic acid or aspartate-semialdehyde in both control and mutant complex I.

This indicated that the oxidative load around the site of ROS production was the markedly higher on glucose media and masked the slight increase in methionine oxidation induced by the mutation that was noted on galactose media (Figure 1E).

### Mutation ND3^S45P^ modulates the A/D transition of complex I

S45 resides within the loop connecting the first two transmembrane helices of subunit ND3 which is part of the ubiquinone reduction pocket (Zickermann *et al*, 2015) and needs to move during catalysis to drive proton pumping (Cabrera-Orefice *et al*, 2018). Nearby C39 is known to be differentially exposed during the A/D transition of complex I (Galkin *et al*, 2008). Changing the serine into a proline is expected to make the loop less flexible. To address the functional consequences of this change, we investigated the effect of mutation ND3^S45P^ on the A/D transition of complex I.

Addition of divalent cations prevents the activation of deactive complex I (Kotlyar *et al*, 1992) and can thus be used to assess the fraction of complex I in the D-form. Mitochondria prepared from cybrid controls that were analyzed in the presence of Mg^2+^ exhibited a rate of NADH oxidation decreased by about 35% of that without this addition. This indicated that the divalent cations prevented activation of about one third of complex I (Figure 1F). In mitochondria prepared from mutant ND3^S45P^, the presence of Mg^2+^ reduced the maximal activity by about 50% indicating that more of the mutant enzyme was in the dormant form. Considering that even in controls, ∼20% of the activity was not inhibitor sensitive (Figure 1D) and thus did not reflect complex I dependent NADH oxidation, the absolute fractions of D-form should be even higher. In any case, an estimated ∼1.5-fold higher fraction of D-form was observed in mutant mitochondria as compared to controls. This suggested that complex I carrying mutation ND3^S45P^ had a higher propensity to deactivate, either already in intact cells or during the preparation of mitochondria. When mitochondria were first deactivated by incubation at 37°C in the absence of substrates, the presence of Mg^2+^ during the experiment reduced the rate in control mitochondria by 70% (Figure 1F). Considering again that most of the residual activity was not due to complex I (Figure 1F, dotted line), this indicated that in controls, almost 90% of complex I were kept locked in the D-form by divalent cations. Mitochondria carrying mutation ND3^S45P^ showed very similar residual activities in the presence of divalent cations, but due to the lower maximal NADH oxidase activities, this represented only a reduction by about 45% (Figure 1F).

To further explore the functional changes induced by mutation ND3^S45P^ at the level of complex I, we measured the temperature dependence of the NADH oxidase activity and the rate of reactivation following complete deactivation. Based on these data, we determined the energy of activation for the transition from the D- to the A-form. For both control and mutant complex I, the relative temperature dependent increase of the catalytic rate was quite similar, amounting to a 1.7 and 1.5-fold increase in the rate of NADH oxidation per 10°C, respectively (Figure 1G). At 25°C, the rate of reactivation was ∼2 min^-1^ for both control and mutant mitochondria, but its temperature dependence was significantly different (Figure 1H). Correspondingly, the energy of activation, as determined from the slope of an Arrhenius plot of these data (Figure 1H), was found to be 61 ± 3kJ·mol^-1^ for control, but only 43 ± 3 kJ·mol^-1^ for mutant complex I (*p*<0.01; n = 3-5). We concluded that, in line with the higher fraction of the D-form in mitochondria as prepared from mutant cells (Figure 1F), mutation ND3^S45P^ rendered complex I more prone to deactivation.

### Mutation ND3^S45P^ moderately affects mitochondrial respiration

To analyze the impact of mutation ND3^S45P^ on the performance of the OXPHOS system, we measured oxygen consumption of isolated mitochondria from control and ND3^S45P^ cybrid cell lines by high-resolution respirometry. The observed reduction in the maximal respiratory rates when feeding electrons via glutamate/malate/pyruvate (Figure 1I) by about 60% was in line with the slight decrease in abundance (Figure 1B) and the markedly lower specific activity (Figure 1C) of complex I. We also noted a ∼20% decrease in succinate-dependent rates (Figure 1J), which seems in line with the observation that complex II abundance trended slightly lower (Figure 1B). As expected for increasing flux through the respiratory chain, the reductions were more pronounced when the mitochondria were in the phosphorylating-active or uncoupled state. Respiratory control ratios, expressed as the ratio of active or uncoupled rate divided by the controlled rate, were significantly reduced only with complex I linked substrates (Figure 1K). This is because oxidizing NADH results in the pumping of 10 protons as compared to just 6 protons per succinate. Importantly, in mutant mitochondria, the respiratory control ratios for complex I linked substrates, albeit lower due to the reduced maximal rates of respiration, were still significantly higher than for the complex II linked substrate (Figure 1K). This indicated that complex I carrying the mutation was still pumping protons. The amount of ATP synthesized per oxygen consumed (P/O ratio) was only slightly reduced with both substrates in mutant mitochondria (Figure 1L). This reflects the expected higher portion of leak-induced losses at lower overall rates. Thus, we concluded that the pumping stoichiometry was not affected by the mutation in complex I, as well. Overall, the observed moderate impairments of OXPHOS function at the level of isolated mitochondria were entirely consistent with the changes caused by mutation ND3^S45P^ at the level of complex I.

### Mutation ND3^S45P^ has no marked effect on the mitochondrial network

Functional deficiency of the OXPHOS system in mammalian cells is often linked to fragmentation of the mitochondrial network. Structured illumination microscopy (Figure 2A**, B**) revealed that this was not the case for the rather moderate functional impairments caused by mutation ND3^S45P^. Cybrids carrying the mutation that were grown on glucose in fact exhibited a slightly more extensive mitochondrial network than control cells. The same was seen as a trend in cells grown on galactose (Figure 2A). Fibroblasts obtained from two independent healthy controls and two patients with heteroplasmy levels of about 80% for mutation ND3^S45P^ exhibited no significant differences in their mitochondrial network that were essentially identical on glucose and galactose. Merely a slight trend to a higher fraction of intermediate network in patient cells grown on galactose was observed (Figure 2B). These findings indicated that even if cells carrying mutation ND3^S45P^ were challenged by an OXPHOS dependent carbon source, this did not result in apparent morphological changes due to respiratory stress in both cybrids and primary patient cells.

**Figure 2:**
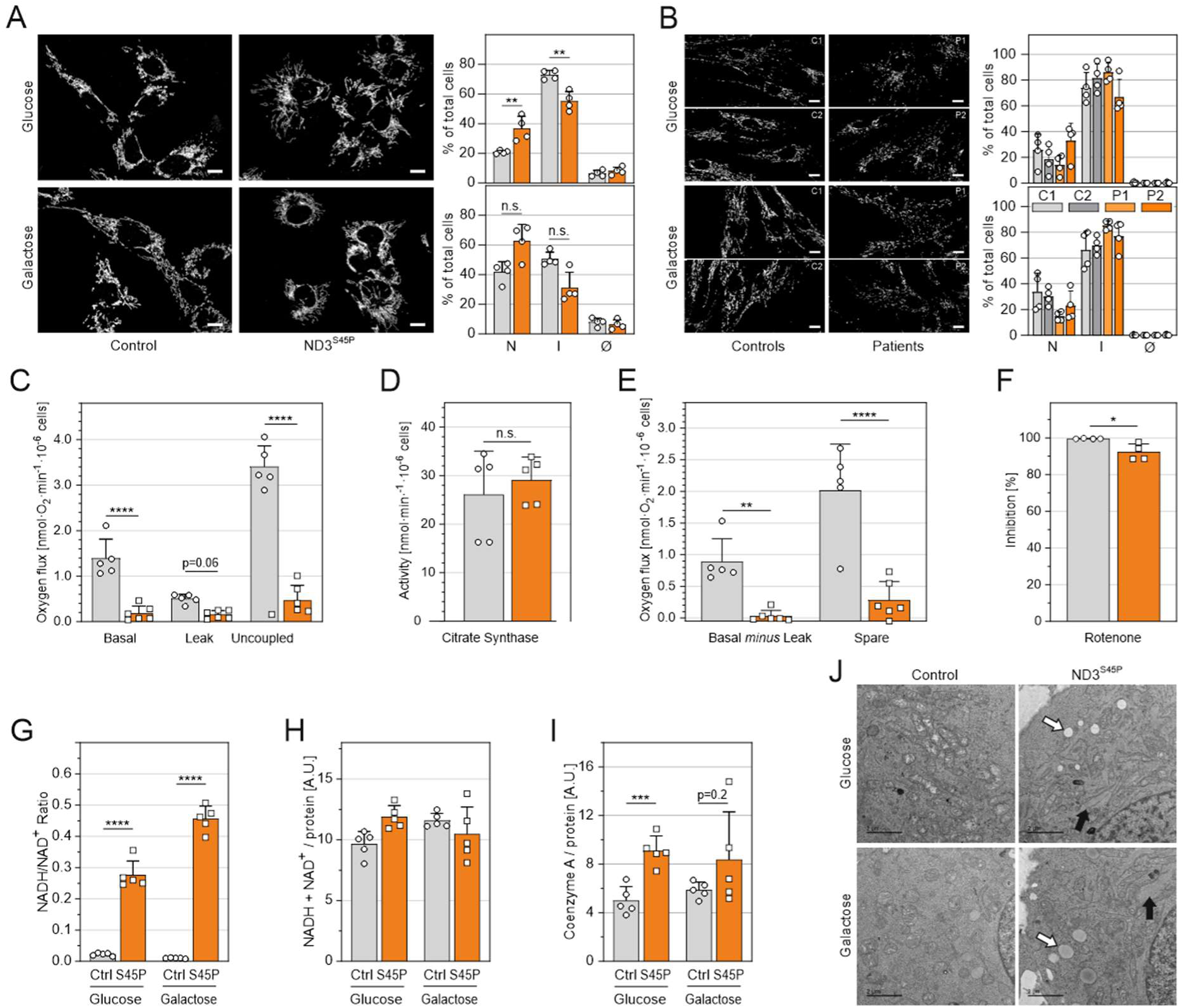
Characterization of intact cells. Mitochondrial Network: The mitochondrial network of control (black/grey) and ND3^S45P^ (orange) cybrids or patient fibroblasts grown on glucose (top) and galactose (bottom) were assessed by structured illumination microscopy using antibodies against the outer membrane protein TOMM20. In each sample, 70-120 cells were classified and the fraction of cells containing an extensive (N), intermediate (I) or no (Ø) network was determined. **A**, control and ND3^S45P^ cybrid cell lines; **B**, controls (C1, C2) and patient fibroblasts (P1, P2). The heteroplasmy levels for mutation ND3^S45P^ in fibroblasts were 75% for patient 1 (P1) and 82% for patient 2 (P2). Whole-cell respiration: Oxygen consumption rates and citrate synthase activities of independently prepared batches of control (grey) and ND3^S45P^ (orange) cybrid cells were measured. **C**, Respiratory rates without additions (Basal), in the presence of complex V inhibitor oligomycin (Leak), and after addition of CCCP (Uncoupled). **D**, citrate synthase activities. **E**, basal minus oligomycin inhibited rates representing basal ATP production (Basal *minus* Leak) and uncoupled minus basal rates indicating spare respiratory capacity (Spare) taken from panel **C**. **F**, inhibition of uncoupled rates by complex I inhibitor rotenone. Measurement of NADH/NAD^+^ and Coenzyme A: The ratios (**G**) and total amounts of NADH and NAD^+^ (**H**) and the amount of Coenzyme A per protein (**I**) were determined by extraction from control (Ctrl, grey) and ND3^S45P^ (S45P, orange) cybrids on glucose and galactose media. A.U., arbitrary units. Electron microscopic analysis: Representative micrographs of control (left) and ND3^S45P^ (right) cybrids on glucose (top) and galactose media (bottom). Mutant cells showing lipid droplets (white arrows) and extended filaments (black arrows) that were not seen in controls are presented. Scale bars 10 µm in **A, B** and 2 µm in **J**. Data are presented as mean ± SD (n = 4-6) and analyzed by Student’s *t*-tests; \**p*<0.05, \*\**p*<0.01, \*\*\**p*<0.001, \*\*\*\**p*<0.0001, n.s. not significant.

### Mutation ND3^S45P^ greatly reduces respiration of intact cells essentially abolishing basal ATP synthesis

Since mutation ND3^S45P^ moderately impaired the functional properties of complex I and mitochon-drial OXPHOS, we next analyzed respiration at the level of intact cells. In contrast to the still rather high oxygen consumption rates of isolated mitochondria, basal and uncoupled respiration of cybrids with mutation ND3^S45P^ was drastically reduced by about 85% (Figure 2C). Unchanged specific activities of citrate synthase (Figure 2D) indicated that this dramatic decrease was not due to reduced amounts of mitochondria per cell. Importantly, the complex V inhibitor oligomycin had almost no effect on the small remaining oxygen flux in mutant cells. This indicated that the residual rate resulted from the mitochondrial proton leak and not ATP synthesis (Figure 2C**, E**). In the presence of the uncoupler carbonyl cyanide *m*-chlorophenyl hydrazine (CCCP), oxygen consumption rates increased about three-fold in both control and mutant cells, showing that there was some spare respiratory capacity in the presence of mutation ND3^S45P^, albeit at a much lower level than would have been expected from the comparably high activities and oxygen consumption rates measured at the level of complex I and isolated mitochondria (Figures 1C**, F, I**). Interestingly, the remaining uncoupled respiratory rate was, if at all, only slightly less sensitive to the complex I inhibitor rotenone (Figure 2F).

### Mutation ND3^S45P^ massively increases the NADH/NAD^+^ ratio and causes accumulation of lipid droplets and filaments in the cytoplasm

To investigate why the respiratory capacity in intact cells carrying mutation ND3^S45P^ did not match the rates of oxygen consumption of mutant mitochondria themselves, we next analyzed the amounts of two coenzymes most critical for driving mitochondrial energy conversion. Strikingly, mutant cells exhibited a massive increase in the NADH/NAD^+^ ratio largely due to ∼10-fold and more than 20-fold increases in NADH on glucose and galactose media, respectively (Figure 2G), while the overall content of NADH plus NAD^+^ was unchanged (Figure 2H). This demonstrated that respiration was not limited by the supply of reducing equivalents. We could also exclude that mitochondrial energy metabolism was impaired by a lack of, for instance, Coenzyme A, which is required to feed substrates into the major catabolic pathways and connect them by transferring acetyl groups between them. On the contrary, the content of Coenzyme A was roughly doubled in mutant cells grown on glucose and trended to be higher on galactose media as well (Figure 2I).

Corroborating the notion that mutation ND3^S45P^ caused a pronounced shift of energy metabolism towards an excess of reducing equivalents, the cytoplasm of mutant cells cultured on both carbon sources was characterized by the presence of many lipid droplets that were hardly observed in the controls (Figure 2J, white arrows). This not only showed that the surplus of reducing equivalents and metabolic substrates was driving fatty acid synthesis resulting in the storage of excess lipids, but also suggested that the mutant cells were not able to sufficiently regenerate NAD^+^ by forming and secreting lactate as typically observed in respiratory impaired cells generating ATP by anaerobic glycolysis.

In line with the still tenable OXPHOS performance in the presence of the mutation (Figure 1I-L) and its limited impact on mitochondrial morphology (Figure 2A**, B**), electron micrographs showed no major effect on cristae morphology (Figure 2J). Interestingly, however, we noted the accumulation of intermediate filaments in cells carrying mitochondrial mutation ND3^S45P^ (Figure 2J, black arrows) further indicating major remodeling at the level of the entire cell.

### Remodeling of the mitochondrial proteome provides insights into metabolic adaptation and derailment

Our observation that NADH oxidation was severely impaired in intact cells but still sustainable in isolated mitochondria suggested that the limited functional deficiencies caused by mutation ND3^S45P^ at the level of complex I induced a misdirected metabolic response exacerbating rather than mitigating the derangement. We reasoned that a broader investigation of the impact of the mutation on the molecular inventory of mitochondria would be required to gain insight into the pathogenic mechanism. Therefore, we used complexome profiling to comprehensively investigate whether the mutation caused specific changes in the mitochondrial proteome that could provide hints to the underlying mechanism causing the metabolic derailment. To explore mitochondrial remodeling in response to respiratory stress, we also forced OXPHOS-mediated aerobic energy conversion by shifting the cells from glucose to galactose as the primary carbon source.

Based on Mitocarta 3.0 (Rath *et al*, 2021), the ∼1000 mitochondrial proteins detected in our complexome profiling dataset represent more than 80% of the total mitochondrial proteome (**Table S1**). For an initial overall assessment of proteome remodeling, we determined the overall impact of mutation ND3^S45P^ and of forcing OXPHOS on the abundance of mitochondrial proteins. Notably, with respect to both, the number and abundance of the proteins affected, the changes exerted by exposing the cells to galactose (a carbon source that does not provide a net yield of ATP through anaerobic glycolysis) were markedly more pronounced than those caused by the respiratory challenge imposed by the pathogenic mutation (**Figure S1**; **Table 1**). Taking control cells on glucose as a reference, the switch to galactose significantly altered the abundance of more than twice as many proteins than introducing mutation ND3^S45P^. Rather than being additive, combining both challenges resulted in abundance changes of just an additional ∼90 proteins. While marked changes in abundance (<0.5x or >2.0x) were found for >60% of the affected proteins by switching from glucose to galactose, this was observed for only ∼30% or less of the proteins affected by the pathogenic mutation. Clearly, mitochondrial remodeling induced by mutation ND3^S45P^ was less pronounced than from exposing the cells to galactose media.

**Table 1:** Number of proteins changed in abundance in complexome profiling datasets.

| Cell Line and Growth Condition | Number of Significantly Changed Proteins ($P < 0.05$) | | |
| --- | --- | --- | --- |
|  | in Total | with Fold Change in Abundance |  |
|  |  | <0.5x | >2.0x |
| Glucose: Control vs. $\text{ND3}^{\text{S45P}}$ | 150 | 28 | 8 |
| Galactose: Control vs. $\text{ND3}^{\text{S45P}}$ | 242 | 44 | 37 |
| Control: Glucose vs. Galactose | 367 | 140 | 101 |
| $\text{ND3}^{\text{S45P}}$: Glucose vs. Galactose | 453 | 148 | 135 |

The four complexome datasets provided a valuable resource to gain molecular insight into the adaptive processes induced by the mutation and the metabolic challenges and how the two responses were connected (**Table S1**).

### Mutation ND3^S45P^ has less effect on markers of mitochondrial ultrastructure, dynamics, quality control and oxidative stress than forcing OXPHOS

We found that the overall abundance of MICOS complex, the multiprotein complex controlling the formation of cristae junctions (Pfanner *et al*, 2014), was reduced by about 60% when cells were grown on galactose instead of glucose. This change was associated with an altered multimeric state of the MICOS complex (Figure 3A). On glucose, mutation ND3^S45P^ caused a reduction in the relative abundances of the core subunits MIC60 and MIC19 by ∼25% that correlated with a reduction of MICOS assemblies larger than 2 MDa.

**Figure 3:**
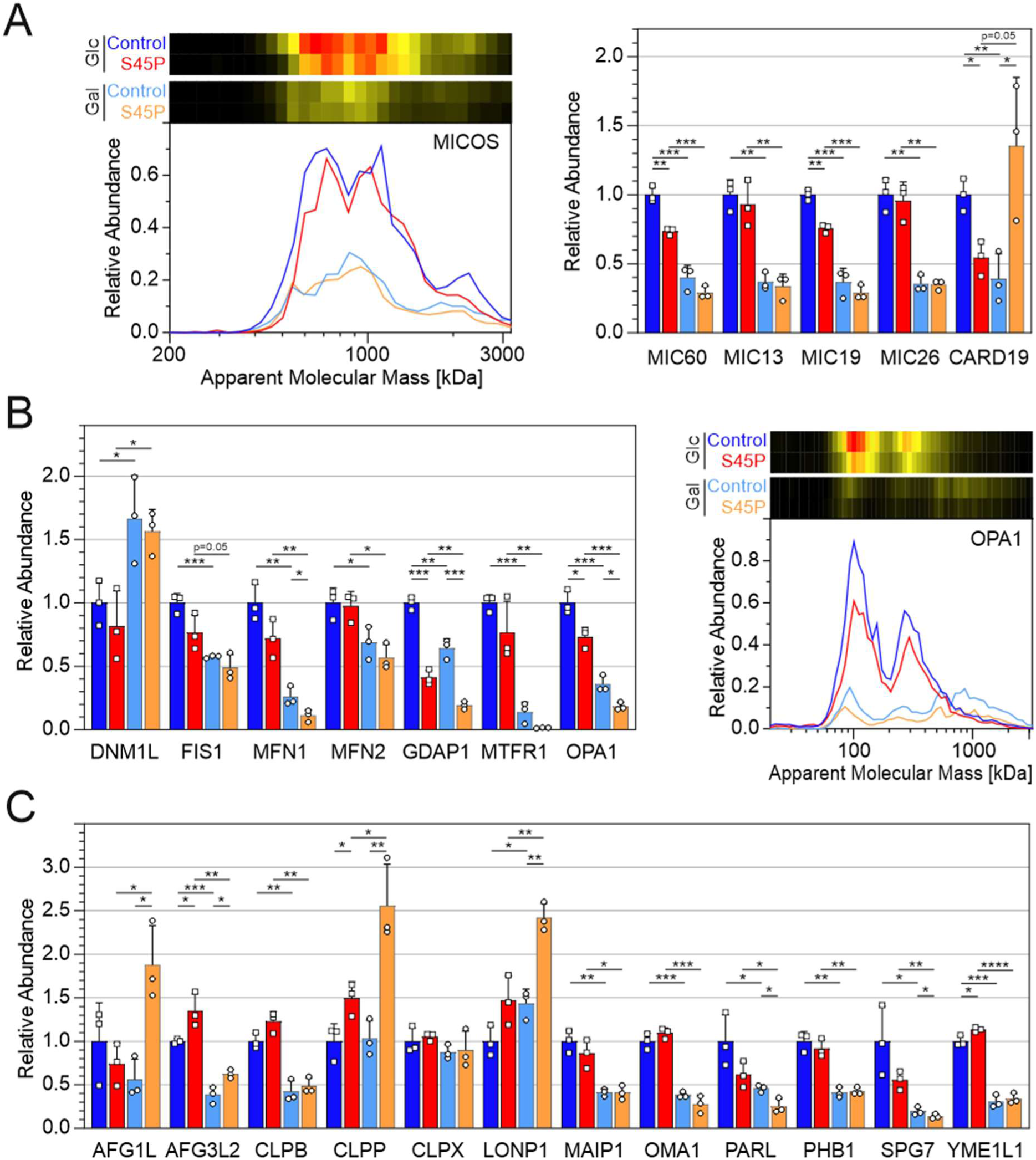
Impact on selected proteins involved in mitochondrial morphology (**A**), dynamics (**B**) and quality control (**C**). Migration profiles and corresponding heatmaps of the MICOS complex (**A**) and OPA1 (**B**) are shown and relative changes for the indicated proteins are presented as bar graphs (**A-C**). Color code: control cells, dark/light blue on glucose/galactose media; cybrids carrying mutant ND3^S45P^, red/orange on glucose/galactose media. Migration profiles and heatmaps in panel A are averages of the profiles for MIC60 (IMMT), MIC13 (CHCHD6) MIC19 (CHCHD6), MIC26 (APOO), and CARD19. Relative abundances were renormalized for each protein by setting the average of controls on glucose to unity and are presented as mean ± SD (n = 3). \**p*<0.05, \*\**p*<0.01, \*\*\**p*<0.001, \*\*\*\**p*<0.0001 (Student’s *t*-test).

In addition, we noted a remarkable pattern of changes in the abundance of CARD19: in control cells on galactose, as well as in ND3^S45P^ cybrids on glucose, CARD19 was decreased by half as compared to controls on glucose. However paradoxically, these levels were restored on galactose media if the mutation was present (Figure 3A). Originally, CARD19 had been proposed to regulate Bcl10-dependent NF-κB activation, but has later been implicated as a factor controlling mitochondrial cristae morphology and recently has been shown to interact with MICOS (Rios *et al*, 2020; Rios *et al*, 2022).

Similarly, OPA1, a protein involved in maintaining cristae morphology, mitochondrial function and mitochondrial dynamics (Del Dotto *et al*, 2017), was somewhat reduced in abundance by two thirds on galactose but was affected much less by the mutation (Figure 3B). Switching the carbon source resulted in marked changes in the migration profiles, most likely reflecting significant changes in the interaction pattern of OPA1 with other mitochondrial complexes, while mutation ND3^S45P^ had only moderate additional effects in further reducing the amounts of larger assemblies.

Galactose-induced changes in the abundance of several proteins involved in controlling mitochondrial dynamics tended to be mildly enhanced by the mutation as well (Figure 3B). However, two regulators of mitochondrial fission were more strongly affected by mutation ND3^S45P^: levels of GDAP1 were reduced by 50% on both media adding to the decrease exerted by galactose alone in control cells. Forcing OXPHOS reduced the abundance of MTFR1 to one fifth and on galactose this protein was hardly detectable at all.

Likewise, several proteases and other proteins involved in mitochondrial quality control were only mildly affected (Figure 3C): if there was any impact on protein abundance by mutation ND3^S45P^, it only slightly reinforced the changes induced by galactose. Only in the case of CLPP, LONP1 and PARL changes were somewhat more pronounced in the presence of the mutation. For AFG3L2, the effects of mutation ND3^S45P^ counteracted the ∼50% decrease caused by the metabolic shift, while for AFG1L an opposing effect of the mutation was evident only on galactose media.

For markers of oxidative defense, a similar pattern of slight enhancement of the changes induced by forcing OXPHOS was evident, because marker proteins like some of the peroxyredoxins, the superoxide dismutases SOD1 and SOD2, as well as thioredoxin-reductase (TXNRD2) tended to respond slightly more to the mutation than to the metabolic challenge (Figure 4A).

**Figure 4:**
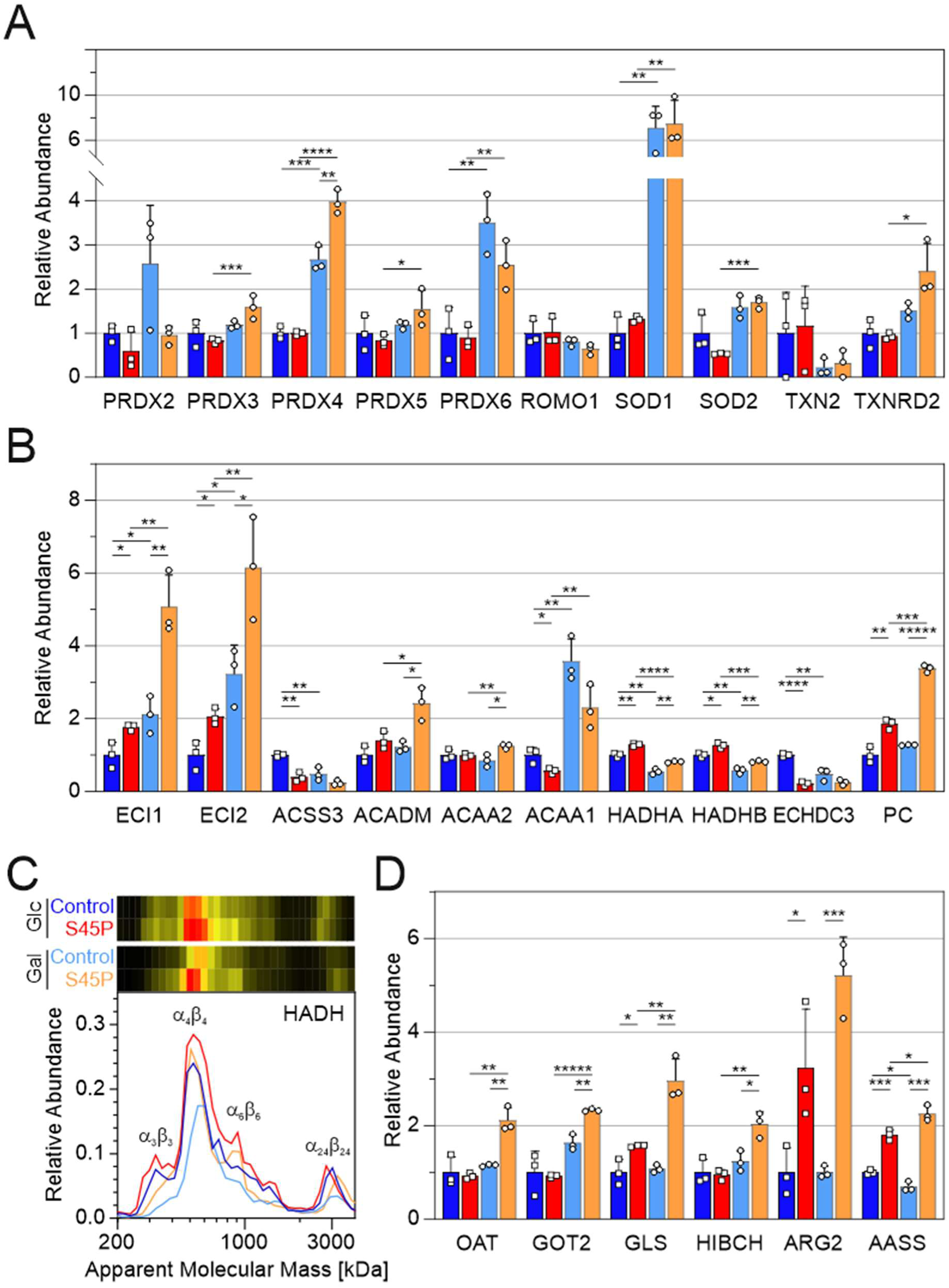
Impact on selected proteins involved in oxidative defense (**A**), β-oxidation (**B, C**) and amino acid metabolism (**D**). **C**, migration profiles and corresponding heatmaps of the α and β subunits of the trifunctional enzyme HADHA and HADHB. Relative changes for the indicated proteins are shown as bar graphs (**A**, **B**, **D**). Color code: control cells, dark/light blue on glucose/galactose media; cybrids carrying mutant ND3^S45P^, red/orange on glucose/galactose media. Relative abundances were renormalized for each protein by setting the average of controls on glucose to unity and are presented as mean ± SD (n = 3). \**p*<0.05, \*\**p*<0.01, \*\*\**p*<0.001, \*\*\*\**p*<0.0001, \*\*\*\*\**p*<0.00001 (Student’s *t*-test).

Since aberrant cristae morphology, mitochondrial fragmentation as well as the activation of quality control systems and oxidative defense mechanisms are hallmarks of OXPHOS deficiency, the limited responses at the level of these systems further support the notion that pathological mutation ND3^S45P^ had rather limited impact on mitochondrial function as such.

### Mutation ND3^S45P^ increases enzymes in some NADH generating pathways partly counteracting metabolic adaptation induced by forced OXPHOS

One way by which mitochondria adapt to maintain proper redox homeostasis is to control the expression level of their catabolic pathways. Therefore, we next analyzed abundance changes of components of the major catabolic pathways in mitochondria as they respond to mutation ND3^S45P^ and to the metabolic challenge by galactose media.

The amounts of several mitochondrial enzymes involved in the metabolism of fatty acids were affected significantly by both, the mutation and forced OXPHOS (Figure 4B): ECI1 and ECI2, two isoforms of enoyl-CoA delta isomerase that feed unsaturated fatty acids into β-oxidation were both found to be increased significantly and in an additive manner by the mutation as well as by switching to galactose. A similar but inverse pattern was observed for ACSS3, the acyl-CoA synthase activating short-chain fatty acids for β-oxidation. ACADM, the medium-chain acyl-CoA dehydrogenase, as well as ACAA2, the major 3-ketoacyl-CoA thiolase, did not respond to a change of carbon source, but in the presence of mutation ND3^S45P^ a significant increase was observed in cells grown on galactose. This contrasts the changes in the abundance of its isoform ACAA1 that is mostly active in peroxisomes but is also found in mitochondria: ACAA1 exhibited a several-fold increase on galactose media, while it was lowered by the complex I mutation. Such opposing effects were most notable for HADHA and HADHB, the α and β subunits of trifunctional enzyme that catalyzes three of the four reactions of the β-oxidation pathway. A reduction in abundance of both proteins caused by forcing OXPHOS was counteracted by an increase caused by mutation ND3^S45P^. On galactose media, this compensatory effect of the mutation amounted to a complete restoration of the major forms of the HADH complex to levels found in control cells on glucose (Figure 4C). Furthermore, we noted two additional, highly significant changes that may have important ramifications for fatty acid homeostasis: ECHDC3 contains an enoyl-CoA hydratase domain and has been implicated in determining insulin sensitivity in humans (Sharma *et al*, 2019). Interestingly, this protein is downregulated almost 5-fold in mitochondria carrying mutation ND3^S45P^, thus exceeding the effect of galactose by far (Figure 4B). Possibly contributing to the formation of lipid droplets in mutant cells (Figure 2J), especially on galactose media, we observed a marked increase of pyruvate carboxylase (PC) known to feed building blocks into fatty acid synthesis (Figure 4B).

Next, we inspected enzymes involved in amino acid catabolism as another important supply of mitochondrial energy metabolism (Figure 4D): While switching from glucose to galactose media had very little effect on control cells, the presence of mutation ND3^S45P^ resulted in marked increases of several key enzymes. Most prominently, OAT and GOT2, two transaminases directly connecting amino acid metabolism to the tricarboxylic acid (TCA) cycle, but also glutaminase (GLS) supplying GOT with glutamate, were increased about two-fold. A similar change in mutant cells on galactose media was evident for HIBCH, an enzyme involved in valine degradation. Furthermore, ARG2, an arginase isoform important for arginine degradation outside of the urea cycle, as well as AASS, a key player in lysine catabolism, were found to be several-fold higher in mutant cells in a fashion largely independent of the carbon source.

Finally, we had a close look at pyruvate dehydrogenase (PDH) and all enzymes of the TCA cycle that supply most of the NADH to the respiratory chain (Figure 5A): Apart from fumarase (FH) and to a lesser extent malate dehydrogenase (MDH2) and aconitase (ACO2), there was no significant effect from forcing OXPHOS in control cells. For mitochondria carrying mutation ND3^S45P^, a mostly slight but significant rise in abundance on galactose media was found for all components of the TCA cycle, except for PDH. Only for aconitase (ACO2) and the GDP-dependent isoform of succinate-CoA ligase (SUCLG2), this specific increase for mutant mitochondria was more pronounced.

**Figure 5:**
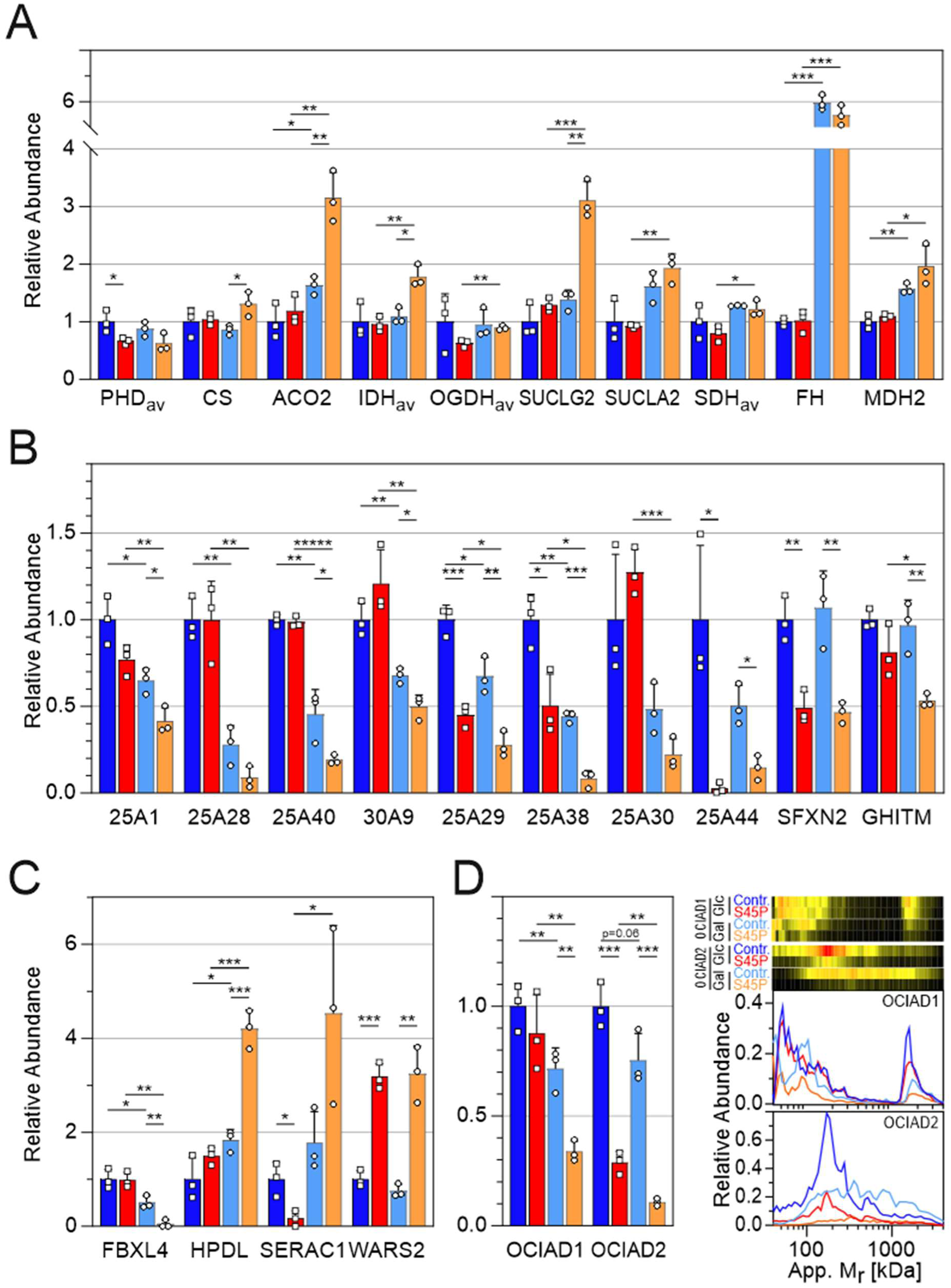
Impact on TCA cycle enzymes (**A**), mitochondrial transporters (**B**) and selected disease-related proteins (**C, D**). **D**, migration profiles and corresponding heatmaps of orphan mitochondrial proteins OCIAD1 and OCIAD2. Relative changes for the indicated proteins are shown as bar graphs (**A-D**). Color code: control cells, dark/light blue on glucose/galactose media; cybrids carrying mutant ND3^S45P^, red/orange on glucose/galactose media. Relative abundances were renormalized for each protein by setting the average of controls on glucose to unity and are presented as mean ± SD (n = 3). \**p*<0.05, \*\**p*<0.01, \*\*\**p*<0.001, \*\*\*\*\**p*<0.00001 (Student’s *t*-test).

### Levels of several mitochondrial carriers are reduced in cells carrying mutation ND3^S45P^

To maintain a proper exchange of metabolites, ions and cofactors with the cytosol, numerous transport proteins are present in the inner mitochondrial membrane. Hence, they play a critical role in maintaining mitochondrial function and homeostasis. Of the many mitochondrial carriers, the abundance of only a limited number was reduced by switching the carbon source or by introducing the mutation (Figure 5B): On glucose media, mutation ND3^S45P^ had no significant effect on the levels of tricarboxylate carrier SLC25A1, iron transporter mitoferrin-2 (SLC25A28), the probable glutathione transporter SLC25A40 and the proton-coupled zinc antiporter (SLC30A9), but it did enhance the decrease seen already in controls when switching from glucose to galactose media. Basic amino acids transporter SLC25A29 and glycine transporter SLC25A38 were lowered markedly in abundance by changing the carbon source and the mutation in an additive fashion. A very pronounced decrease by more than five-fold was also observed for the inorganic ion carrier SLC25A30 on galactose, but only in the presence of mutation ND3^S45P^. A massive impact of the mutation was seen for branched-chain amino acids transporter SLC25A44 that was reduced to levels close to the detection limit on both carbon sources. Also, SFXN2, one of several isoforms of the serine transporter family of sideroflexins, was about halved by the mutation on glucose and galactose as compared to control that showed no effect from forced OXPHOS. Finally, the calcium/potassium to proton antiporter GHITM that is assumed to play a regulatory role in mitochondrial maintenance (Patron *et al*, 2022) caught our attention, because on galactose, mutation ND3^S45P^ significantly reduced its abundance about two-fold.

### Several other proteins implicated in mitochondrial disorders are affected by mutation ND3^S45P^

We noted significant changes in the abundance of a number of proteins linked to mitochondrial diseases (**Table S1**) of which six especially remarkable examples will be briefly highlighted here (Figure 5C**, D**). Deficiency of FBXL4, a component of E3 ubiquitin ligase complexes (Skaar *et al*, 2013), causes mtDNA depletion syndrome (Bonnen *et al*, 2013) and promotes mitophagy (Alsina *et al*, 2020). On glucose, mutation ND3^S45P^ had no effect on FBXL4 levels, but it aggravated a reduction in abundance induced by galactose rendering it almost undetectable. Mutations in HPDL, a dioxygenase involved in the metabolism of aromatic amino acids, lead to early onset neurodegenerative disorders (Husain *et al*, 2020). On galactose but not on glucose, mutation ND3^S45P^ consistently more than doubled the level of HPDL with high significance. SERAC1 is a phospholipid remodeling enzyme and its deficiency causes Leigh-like syndrome (Wortmann *et al*, 2012). We found that in control cells SERAC1 levels were hardly affected by forcing metabolism into OXPHOS dependence, but switching the carbon source for cells carrying mutation ND3^S45P^ changed SERAC1 abundance levels substantially and into the opposite direction than the mutation did in cells on glucose media. While in mutant cells SERAC1 abundance was less than one fifth of controls on glucose, it was more than doubled on galactose thereby showing a pattern similar to the one described above for CARD19. Mutations in the mitochondrial tryptophan-tRNA-synthase WARS2 cause neurodevelopmental disorders (Wortmann *et al*, 2017), and mutation ND3^S45P^ consistently increased its abundance by more than three-fold independent of the carbon source (Figure 5C).

Finally, OCIAD1 and OCIAD2, named after their ovarian carcinoma immunoreactive antigen-like (OCIA) domain, caught our attention. The two orphan mitochondrial proteins were reported to be associated with amyloid β production and neuronal vulnerability in Alzheimer’s disease (Han *et al*, 2014; Li *et al*, 2020). Both proteins were implicated to play a role in STAT3 signaling, but more recently it has been proposed that they play a role in complex III assembly (Chojnacka *et al*, 2022; Le Vasseur *et al*, 2021). Under the conditions used here, OCIAD1 and OCIAD2 for the most part seemed not to be associated in the same complex. Instead, we found that both independently exhibited pronounced variability in their multimeric states depending on the carbon source. However, the stability of the observed complexes seemed only marginally affected by mutation ND3^S45P^, although its presence did lower OCIAD1/2 levels several-fold overall (Figure 5D). On glucose, OCIAD1 mostly migrated over a broad range of 50-200 kDa apparent mass. Still, a substantial amount of the protein was also found in large complexes migrating at around 1.5, 2.0 and 3.0 MDa. On galactose, the 1.5 MDa complex but not the larger forms disappeared and at lower masses more defined peaks at about 40-50 kDa and 90-100 kDa became visible. In contrast, OCIAD2 showed fairly defined peaks around 170 kDa and to a lesser extent at ∼500 kDa that completely disappeared on galactose media giving way to a broad distribution of the protein at 100 kDa and above, all the way to apparent masses of several MDa. Since we could not find other proteins following a similar migration pattern, it seems that the OCIAD1 and OCIAD2 complexes at different masses represent multimeric states of the proteins. Mutation ND3^S45P^ seemed not to affect the remarkable plasticity of the two OCIA-domain proteins. However, in the presence of mutant ND3^S45P^, OCIAD1 was reduced by more than 50% on galactose, but not on glucose, while compared to controls OCIAD2 levels were decreased several-fold on both carbon sources.

### Analysis of the impact of mutation ND3^S45P^ in patient fibroblasts confirms the pattern of moderate mitochondrial remodeling

To track the impact of mutation ND3^S45P^ all the way from the molecular level of complex I function to the cellular level, we performed the analysis in cybrids, since they are homoplasmic for the mutation and enough mitochondria can be obtained. However, cybrids are derived from tumor cell lines that can potentially adapt to the pathogenic challenge over multiple division cycles. This is of less concern for the biochemical analysis of the affected enzyme complex itself but may significantly affect the analysis at the level of entire mitochondria and cells. If so, analyzing the impact on cybrid mitochondria may still provide important insights into the remodeling mechanisms caused by the molecular defect, but caution needs to be applied when transferring such findings to the situation in patient cells. Therefore, we went on to analyze complexomes of mitochondria from fibroblasts of two patients with a heteroplasmy level of about 80% for mutation ND3^S45P^ in comparison to two healthy controls (**Table S2**). We again tested the impact of shifting these fibroblasts from glucose to galactose media (**Table S1**). Since data for samples from just two patients were available, a statistical analysis was not feasible. However, consistent trends in mitochondrial remodeling in cells from different individuals still allowed us to assess whether the changes observed in cybrids were relevant overall in the context of primary cells from patients as well.

Changes in the mitochondrial proteome exerted by shifting from glucose to galactose generally seemed to be less pronounced and affected a markedly lower number of proteins in fibroblasts than in cybrids. This most likely reflected a larger capacity of the highly proliferative tumor cells to adapt to metabolic challenges (**Table S1**). Nevertheless, also in fibroblasts, the impact of switching the carbon source clearly seemed to exceed that of introducing mutation ND3^S45P^ alone (**Figure S2**). Importantly, the expression of OXPHOS complexes was hardly affected and changes in markers of mitochondrial stress induced by both forcing OXPHOS and the mutation tended to be less prominent in fibroblast mitochondria as well.

To explore to what extent the pattern of mitochondrial remodeling seen in cybrids could also be observed in fibroblasts and to look out for specific differences, we had a closer look at some of the complexes and proteins that we had focused on in cybrids (Figures 3-5).

For MICOS, the organizer of mitochondrial morphology, fibroblasts exhibited a much higher propensity to form larger complexes on glucose, while the pattern of complexes was very similar for both cell types on galactose (**Figure S2A**). As in cybrids, switching the carbon source reduced the overall abundance of MICOS components by about 50% and mutation ND3^S45P^ tended to have very little effect in fibroblasts. The putative MICOS interactor CARD19 exhibited a similar tendency to be increased by the mutation on galactose in fibroblasts, while there were no indications for the decrease observed in cybrids on glucose media. The latter was also seen for OPA1 with the effects of both carbon source and mutation seeming much less pronounced overall (**Figure S2A**). As for MICOS, the pattern of association of OPA1 with different complexes was much more affected in cybrids than in fibroblasts and largely independent of whether mutation ND3^S45P^ was present or not. Notably, switching from glucose to galactose seemed to selectively reduce the association of OPA1 with complexes larger than ∼700 kDa (**Figure S2A**).

Likewise, the abundance of proteins involved in mitochondrial dynamics was less affected by forcing OXPHOS in fibroblast than in cybrid mitochondria and in many cases, no clear trend was evident (**Table S1**). Of the two proteins that we had identified as being more affected by mutation ND3^S45P^, GDAP1 showed a similar abundance pattern in both cell types, whereas for mitochondrial fission regulator MTFR1, the effects of the mutation seemed to be inverted (**Figure S2B**). While its amount clearly trended to be strongly reduced in fibroblasts but not in cybrids on glucose, it seemed to reach glucose control levels in the patient cells rather than being almost undetectable in ND3^S45P^ cybrids (Figure 2B). Also, for proteins involved in mitochondrial quality control, the already small effects of mutation ND3^S45P^ seen in cybrids for CLPP, LONP1 and PARL seemed to be absent in fibroblasts. Except for PARL, no effect of switching from glucose to galactose media was observed in fibroblasts (**Figure S2B**; **Table S1)**. For markers of oxidative defense, hardly any effects from the mutation or from switching the carbon source were seen in fibroblasts. PRDX6 and SOD1 even tended lower rather than showing marked and significant increases as in cybrid mitochondria upon forcing OXPHOS (Figure 4A**, S2B**).

As illustrated for several examples (**Figure S2B, C**), the marked changes in abundance induced by mutation ND3^S45P^ - as observed in cybrid mitochondria for some enzymes of β-oxidation, amino acid metabolism and TCA cycle (Figures 4B**, D; 5A**) - were not seen in fibroblasts (**Table S1**, **Figure S2B**). If anything, the abundances of these proteins tended to opposite directions when comparing the two cell types. The same was true for the mitochondrial transporters, since even for the prominent mutation-induced changes in cybrids for SLC25A38, SLC25A44 and SFXN2, there were no indications for abundance changes in fibroblast mitochondria. The only exception was the inorganic iron carrier SLC25A30 which, on galactose, decreased similarly in both cell types (**Figure S2C**).

Of the six disease-related proteins that changed significantly in response to mutation ND3^S45P^ in cybrids (Figure 5C**, D**), hardly any effects were visible for HPDL, SERAC1, WARS2 and OCIAD1 in fibroblast mitochondria (**Figure S2C; Table S1)**. However, the pattern of abundance changes seemed similar in both cell types for OCIAD2. Contrary to the general trend, the amounts of FBXL4 appeared to be even more drastically reduced by the mutation and for forced OXPHOS in fibroblasts (**Figure S2C**).

In summary, the reduced abundance changes for the proteins in focus in the context of this study reflected the overall trend seen for the mitochondrial proteome. Still, a number of apparent similarities and notable differences in the pattern of mitochondria between the two types of cells were observed.

### Analysis of the impact of mutation ND3^S45P^ in patient muscle tissues suggests cell type specific differences in mitochondrial remodeling

While fibroblasts are widely used to assess mitochondrial disease-related changes at the cellular level, the impact on patient tissues may still be quite different. As the only human tissue available from patients, we thus performed complexome profiling on muscle biopsies from the same two patient donors of the fibroblasts, to analyze and compare it to muscle tissue from three healthy controls (**Table S1**). The heteroplasmy levels in the patient muscle tissues were confirmed to be ∼80% (**Table S2**). Again, due to the limited number of patients available, only trends could be observed.

Variants of the complexes containing MICOS components with larger masses were more prominent in muscle tissues than in cybrids and patient fibroblasts, but like in cells, mutation ND3^S45P^ seemed to have no marked effect on the stability and abundance of the complexes (**Figure S3A**). Notably, in contrast to fibroblasts (**Table S1**), the abundances of the markers of mitochondrial quality control AFG3L2, MAIP1, OMA1 trended lower in muscle from patients as compared to healthy controls (**Figure S3B**). However, there were no indications for the prominent increases in the markers of oxidative defense SOD1, SOD2 and PRDX6 in muscle mitochondria (**Figure S3B**) that we observed in cybrid mitochondria (Figure 4A).

Likewise, for some of the metabolic enzymes, a tendency for mutation-induced abundance changes was seen, that was not evident in fibroblasts from patients, but resembled the situation in cybrids (**Figure S3C**): the amounts of ACADM and both subunits of the trifunctional enzyme HADH of β-oxidation and ACO2 and SUCLG2 of TCA cycle trended higher, while the abundance of the glycine transporter SLC25A38 seemed to be reduced in patient muscle samples. Notably, the size distribution of HADH oligomers was not affected by the mutation (**Figure S3D**). Of the six disease-related proteins that were significantly affected by mutation ND3^S45P^ in cybrids (Figure 5C**, D**), only OCIAD1 exhibited a slight trend for lower amounts in patient muscle as well (**Figure S3C**).

As for patient fibroblast mitochondria, mutation ND3^S45P^ seemed to lead to less extensive mitochondrial remodeling in patient muscle tissue than in cybrids overall. Interestingly however, several mutant-induced changes seen in cybrids seemed to occur in muscle biopsies but not in fibroblasts from the same individuals, pointing towards a cell type- and tissue-specific response to the mutation at the level of the mitochondrial proteome.

## Discussion

To gain insight into the still poorly understood pathogenic mechanisms of inherited mitochondrial disorders, we explored some of the uncharted territory between a severe disease-causing mitochondrial mutation and its often heterogeneous and variable patient phenotypes. We took the actual molecular defect as a starting point for a stepwise and in-depth analysis of its impacts on the affected enzyme, its mechanism of catalysis, and went on with evaluating mitochondrial function, integrity and proteomic remodeling. Mutation m.10191T>C was chosen because it directly affects the catalytic core of mitochondrial complex I and this mutation is associated with a rather well-known mtDNA-linked mitochondrial disorder that causes Leigh syndrome, a very severe condition frequently leading to early death (Chang *et al*, 2020; Taylor *et al*, 2001). In addition, this mutation is a typical example of the discrepancy between still rather high activities of the affected OXPHOS complex and a nevertheless devastating clinical outcome.

Point mutation m.10191T>C locates in the mitochondrial gene encoding subunit ND3 of respiratory complex I and causes an amino acid exchange at position 45 that resides in the loop connecting the first two transmembrane helices (loop TMH1-2^ND3^). Subunit ND3 is located in the membrane arm (P module) of complex I and the affected loop is part of its connection to the peripheral arm reaching into the ubiquinone reduction pocket of the Q module (Zickermann *et al*, 2015). Loop TMH1-2^ND3^ harbors the unique cysteine (C39 in humans) that is accessible to chemical modification in the D-form but not the A-form of the enzyme in the Active/Deactive (A/D) transition (Galkin *et al*, 2008). This loop is critical for the mechanism of complex I function because preventing it from moving during the catalytic cycle abolishes proton pumping without affecting ubiquinone reduction (Cabrera-Orefice *et al*, 2018). An amino acid exchange in a mobile domain is unlikely to have major effects on the structural integrity of the multiprotein complex. However, replacing a serine by a proline is expected to render loop TMH1-2^ND3^ less flexible thereby providing a straightforward rationale, on how mutation ND3^S45P^ could interfere with complex I function and regulation.

Indeed, we found that the stability and assembly of mutated complex I were just mildly affected, while the impact on all tested functional properties was notable (Figure 1). NADH oxidase activity sensitive to two complex I inhibitors that bind to overlapping but distinct sites within the ubiquinone binding pocket (Zickermann *et al*, 2008) was reduced by ∼70% in mutant mitochondria. Reflecting limited structural changes in the binding pocket, the affinity of the inhibitors was found to be only slightly different in mutant ND3^S45P^. Interestingly, the A/D transition of complex I was clearly affected by mutation ND3^S45P^, lowering the energetic barrier for this regulatory interconversion by ∼20 kJ·mol^-1^ and resulting in a higher propensity for the D-form.

In good agreement with the degree of complex I deficiency, mutation ND3^S45P^ reduced NADH-dependent oxygen consumption of intact mitochondria by about 60% in the active and uncoupled states and resulted in a corresponding decrease in respiratory control ratios (Figure 1I**, K**). When complex I was circumvented by feeding electrons into the respiratory chain through the succinate dehydrogenase (complex II), oxygen fluxes were still reduced by 20-30% (Figure 1J) serving as a first indication that mutation ND3^S45P^ affected mitochondrial metabolic capacity indirectly as well.

Importantly, P/O ratios were reduced by just ∼15% in NADH- and succinate-dependent respiration (Figure 1L) indicating that the efficiency of ATP synthesized by the OXPHOS system per oxygen consumed to oxidize NADH was only slightly affected by the complex I deficiency. This was due, in other words, to a generally reduced efficiency of energy conversion at lower oxygen fluxes rather than a functional change of complex I itself. In fact, since through the chemiosmotic mechanism of OXPHOS, respiration and ATP synthesis are strictly coupled to the number of protons pumped across the inner mitochondrial membrane per oxygen consumed, the almost unchanged P/O ratio with NADH-dependent substrates showed that mutation ND3^S45P^ had not changed the proton pumping stoichiometry of complex I.

Overall, the extent of complex I deficiency and the defined functional changes at the level of mitochondria seemed consistent with the structural change exerted by mutation ND3^S45P^ on complex I at a region critical for its energy converting mechanism and regulation. Yet, given that complex I still retained about one third of its maximal NADH oxidation and proton pumping capacity, respiratory deficiency alone seemed not sufficient to explain why the mutation leads to such a devastating patient phenotype.

Oxidative stress is commonly considered as a major contributing factor in the pathogenic mechanism of mitochondrial diseases and has been implicated to play an important role in Leigh syndrome models (Adjobo-Hermans *et al*, 2020; Kahlhöfer *et al*, 2017). However, we found that mutation ND3^S45P^ seemed to have only a limited effect on the oxidative load in mitochondria. Although we detected a 10-20% increase in the oxidation of sensitive methionines near the site of ROS production of complex I, this was only seen when oxidative metabolism was forced by shifting cells to galactose media. This notion was corroborated by the finding that the mutation itself had only minor effects on the levels of markers of the antioxidant defense systems (Figure 4A). Notably, however, switching from glucose to galactose as the carbon source resulted in a marked, in some cases several-fold increase in the amounts of several peroxiredoxins and both mitochondrial superoxide dismutases. In fact, this observation showed that mitochondria responded to forcing OXPHOS by ramping up their oxidative defense, which was nicely mirrored in a decreased local oxidative load on galactose as evident from lower levels of methionine oxidation in subunit NDUFS1 as compared to glucose (Figure 1E). In any case, while the reduced complex I activity in the mutant clearly resulted in somewhat higher ROS production rates, it appeared unlikely to be a major contributor to disease severity, because just growing cells on glucose media alone seemed to lead to an equally high or even higher oxidative load.

Indeed, mitochondria from cybrids and fibroblasts, derived from the skin of patients carrying a high mutation load, exhibited hardly any of the characteristic changes of OXPHOS deficiency in mitochondrial dynamics and morphology (Figure 2A**, B, J**) or in the abundance or multimeric state of the proteins controlling them (Figures 3A**, B; S2A, B**). Accordingly, the very limited impact on the levels of markers of mitochondrial quality control in cybrids (Figure 3C) as well as fibroblasts (**Figure S2B**) and muscle tissue (**Figure S3B**) from patients seemed indicative of rather low levels of mitochondrial stress. Notably, remodeling of the mitochondrial proteome in general (**Figure S1; Table 1; S1**) that resulted from shifting cybrids and fibroblasts to galactose to force OXPHOS-dependent metabolism was clearly more pronounced than the changes exerted by mutation ND3^S45P^ (Figures 3**, 4; S2**).

In stark contrast to the comparably moderate impact on parameters of mitochondrial function and oxidative stress, we found that intact cells carrying mutation ND3^S45P^ in cybrids were severely impaired in their capacity to oxidize NADH. This was evident from a ∼85% reduced whole-cell respiration (Figure 2C-F) and a massive increase in the NADH/NAD^+^ ratios on glucose and galactose media (Figure 2G). Notably, these profound differences were not due to lower overall contents of the co-substrate (Figure 2H) or mitochondria (Figure 2A**, B, D**). Accumulation of lipid droplets in the cytoplasm of ND3^S45P^ cybrids served as another striking indication of a massive excess of reducing equivalents in mutant cells that seemingly could not be compensated by increased anaerobic glycolysis, even on glucose media (Figure 2J). While such severe metabolic derailment provided a plausible explanation for the fatal patient phenotypes, it seemed surprising that the quite sizable remaining OXPHOS capacity was not sufficient to maintain low NADH/NAD^+^ ratios in the presence of mutation ND3^S45P^.

We concluded that it was not sufficient to consider just the immediate consequences on complex I activity and mitochondrial energy supply or ROS production in order to understand the pathogenic mechanism of mutation ND3^S45P^. Since the deleterious accumulation of NADH was only observed in the context of intact cells, we deemed it necessary to take a broader perspective by considering derailed adaptive mechanisms and mitochondrial remodeling as well. Nevertheless, we reasoned that any such pathogenic mechanism at the whole cell level should be connected to functional changes of complex I that was found to almost completely control mitochondrial oxygen consumption and that was almost completely stopped by rotenone in both control and mutant cells (Figure 2F).

The reduced activation barrier for the A/D transition and the high propensity of complex I for the deactive state (Figure 1G**, H**) were the only clues from our functional analysis of mitochondria from ND3^S45^ mutant cells providing a possible regulatory link by which mitochondrial NADH consumption could be switched off. Therefore, we hypothesize that lowering the critical threshold of the A/D transition by mutation ND3^S45^ may have led to an overshooting response and aberrantly shutting off mitochondrial respiration and exacerbating the pathological impact. Thereby, cells would become unable to utilize their substantial remaining OXPHOS capacity, in turn providing a reasonable explanation for the extremely severe patient phenotypes caused by the mutation. Under normal conditions, switching complex I into the deactive state may provide a mechanism by which cells are able to adjust mitochondrial respiration to their metabolic needs. This would assign a previously unknown regulatory role to the A/D transition of complex I that has so far been implicated predominatly in the context of ischemia/reperfusion injury (Chouchani *et al*, 2016; Dröse *et al*, 2016).

While a deviant regulation of complex I may offer a straightforward rationale behind the incisive patient phenotypes caused by mutation ND3^S45^, other adverse changes in mitochondrial metabolism may have contributed to the massive accumulation of NADH in the mutant cells. Specifically, we considered that a surge of the NADH generating pathways or impairment of metabolite transport between cytosol and mitochondria may have overloaded the reduced respiratory capacity in mutant cells as well. Indeed, we did observe that mutation ND3^S45^ caused some uptick in the abundance of several components of the TCA cycle and fatty acid oxidation pathways, but often these changes were small and only significant on galactose media (Figures 4B, **5A**; **Table S1**). Again, shifting from glucose to galactose had more pronounced effects on the abundance of some enzymes in these pathways than the mutation. With the notable exception of key enzymes involved in the degradation of basic amino acids and glutamine, a similar pattern of remodeling was observed for proteins involved in amino acid catabolism (Figure 4D; **Table S1**). Interestingly, basic amino acid transporter SLC25A29, glycine transporter SLC25A38, branched-chain amino acid transporter SLC25A44 and one of the isoforms of the serine-transporting sideroflexins (SFXN2) were the only metabolite transporters exhibiting a marked and consistent decrease in mutant cells that significantly exceeded the attenuation seen on galactose media (Figure 5B). Importantly, except for a moderate increase in OAT and GOT2 on galactose only, no mutant-induced changes in the amounts of the mitochondrial carriers and transaminases involved in the aspartate/glutamate shuttle or α-glycerophosphate dehydrogenase were detected (Figure 4D; **Table S1**). Similar trends for these pathways were observed in fibroblasts and muscle tissue from patients. However, changes tended to be even less pronounced and to affect fewer enzymes (**Figures S2B, C; S3C**). Overall, we concluded that the limited increases in NADH generating and shuttling pathways may have contributed to the increased NADH levels especially on galactose media but could not explain the stalled respiration in mutant cells.

In conclusion, our analysis of complex I mutation ND3^S45^ shows that the pathogenic mechanism leading from a range of limited functional impairments at the level of complex I to the most severe patient phenotype of Leigh syndrome is a multifaceted process leading to a failed metabolic response and a derailment of cellular homeostasis. Ultimately, massive accumulation of NADH appeared as the main culprit responsible for the fatal consequences of the mutation that seemed to be connected to an aberrant threshold for the A/D transition of complex I and pointed at a physiological control of complex I activity beyond the NADH/NAD^+^ ratio itself and classic mechanisms of respiratory control. Nothing is known about the signals and factors involved in such a hypothetical regulatory mechanism. In search for the possible players involved, we identified some remarkable changes in the abundance of several proteins that were previously implicated in mitochondrial disorders (Figure 5C**, D**), but it remains obscure how they could be connected to complex I regulation. However, the comprehensive complexome profiling datasets of mitochondria from cybrids and fibroblasts from patients obtained under two metabolic regimes that are complemented by matching data from muscle biopsies (**Table S1**) provide a rich resource for further studies of the mitochondrial remodeling caused by mutation ND3^S45^ and the mechanisms of aberrant regulation of complex I resulting from it.

In general terms, our study is a prime example of how a known and seemingly simple defect caused by a mutation in a well-studied mitochondrial complex may lead to unexpected consequences that go far beyond the immediate functional deficiencies of the affected protein itself. Therefore, much more in-depth analyses of different pathogenic mutations will be needed to solve the conundrum of genotype-phenotype relationships in mitochondrial disease. Importantly, our finding, that aberrant regulation of complex I may severely exacerbate the physiological consequences and thus the disease phenotype, could serve as a starting point for novel therapeutic strategies to tackle mitochondrial disorders.

## Materials and Methods

### Patient samples

Primary forearm skin fibroblasts and muscle (musculus quadriceps or musculus semitendinosus) biopsies (needle or open biopsies) were available from two unrelated patients carrying mutation m.10191T>C and from three unrelated control individuals (**Table S2**), whereby no fibroblast specimen was available from the control 3 subject. The muscle samples were snap-frozen after collection and stored at −80°C. The study was performed in accordance with the Declaration of Helsinki, and informed consent was obtained prior to inclusion in this study.

### Cell culture

Human osteosarcoma 143B cell line (ATCC) used as control and mutation m.10191T>C (ND3^S45P^) of trans-mitochondrial cytoplasmic hybrids (cybrids) with a 143B background (Radboud Biobank) were placed on culture dishes in Dulbeccós modified Eaglés medium (DMEM) containing 4.5 g/l glucose, without L-glutamine and sodium pyruvate(Lonza) supplemented with 10% (v/v) heat-inactivated fetal bovine serum (FBS; Gibco), 1% non-essential amino acids, 1 mM sodium pyruvate, 2 mM L-glutamine, and 50 µg/ml uridine, 100 units/ml penicillin, 0.1 mg/ml streptomycin (all from Sigma). Cell cultures were maintained in a humidified atmosphere of 95% air/5% CO_2_ at 37°C. Before reaching confluence, cells were harvested by trypsinization (0.05% trypsin, 0.02% EDTA; Gibco).

Human fibroblasts were obtained from skin biopsies from two controls and two patients with a heteroplasmy level for m.10191T>C of 75% (P1) and 82% (P2). Fibroblasts were cultured until grown to confluency in M199 medium with Earle’s Balanced Salt Solution (EBSS), L-glutamine, and 2.2 g/l NaHCO_3_ (PAN Biotech) supplemented with 10% (v/v) FBS (Gibco) and 1% penicillin/streptomycin antibiotics (100 U/ml and 0.1 mg/ml, respectively) at 37°C in a humidified atmosphere of 95% air/5% CO_2_. Primary human skin fibroblasts were collected by trypsinization using trypsin powder 1:250 without EDTA (Gibco).

To culture both fibroblast and cybrid cell lines in the presence of galactose, a basal DMEM medium containing only 4 mM L-glutamine but deprived of glucose and sodium pyruvate (Gibco) was used. This medium was supplemented with 5 mM D-galactose (Sigma) and 10% (v/v) dialyzed fetal bovine serum (Gibco), plus 50 µg/ml uridine and 100 U/ml penicillin, 0.1 mg/ml streptomycin. Cells were harvested by trypsinization and stored, if necessary, at –80°C until use.

### Isolation of mitochondria from cybrids and fibroblasts

Cultured cybrids or fibroblasts were harvested by trypsinization, resuspended in their respective growth medium and pelleted by centrifugation at 700 x *g* for 5 min. After washing with cold PBS (pH 7.2), cells were resuspended in ice-cold isotonic isolation buffer (250 mM sucrose, 1 mM EDTA, 20 mM, Tris-HCl, pH 7.4) supplemented with 1 mg/ml fatty acid-free bovine serum albumin (BSA), if mitochondria were to be used for functional tests, or with 1x protease inhibitor cocktail (SigmaFAST), if mitochondria were to be used for complexome profiling analyses. Cells were disrupted mechanically with 12-17 gentle strokes (1800 rpm) using a motor-driven, tightly fitted 5 ml glass/Teflon Potter-Elvehjem homogenizer at 0°C. Mitochondria and mitochondrial membranes were isolated by stepwise differential centrifugation. Intact cells, their debris and nuclei were removed from the homogenate by low-speed centrifugation at 1,000 x *g* for 10 min at 4°C. The resulting crude mitochondrial fraction was harvested from the supernatant by sedimentation at 10,000 x *g* for 10 min at 4°C. For some experiments, crude mitochondrial fractions were further purified by resuspension in isolation buffer and loading on a two-layer (34%/50% (m/v)) sucrose gradient with 1 mM EDTA, 20 mM Tris-HCl, pH 7.4. After centrifugation at 60,000 x *g* for 20 min at 4°C in a swing-out rotor (MLS-50, Optima Max-XP, Beckman Coulter), the mitochondrial fraction localized in the interphase between the two sucrose layers was carefully removed with a syringe, pooled and washed once with isolation buffer (without BSA). Finally, pure mitochondria were sedimented at 22,000 x *g* for 10 min at 4°C and resuspended in 200-400 µl of isolation buffer.

Protein concentration was determined using the Lowry method (DC protein assay kit; Bio-Rad). Fatty acid-free BSA was used as standard. Mitochondrial membranes were snap-frozen in liquid nitrogen and stored at −80°C until use. For analysis of OXPHOS function, intact and coupled mitochondria were used, which is why the crude fractions were prepared from freshly harvested cells and used immediately.

### Isolation of mitochondria from human muscle

Human muscle biopsies were treated as described (Janssen *et al*, 2006). In brief, muscle tissue was resuspended and homogenized in 250 mM sucrose, 1 mM EDTA 20 mM Tris-HCl, pH 7.4 supplemented with 1x protease inhibitor cocktail (SigmaFAST). The homogenate was centrifugated at 1,000 x *g* for 10 min at 4°C; this step was repeated 2-3 times. The resulting supernatant was then centrifuged at 22,000 x *g* for 10 min at 4°C to obtain the crude mitochondrial fraction. Due to limitations in biopsy materials, the supernatant was concentrated again using Amicon 3K ultrafiltration units (Millipore) and was pooled into the crude mitochondrial fraction. Protein concentrations were measured using the Lowry method.

### Activity measurements

To ensure accessibility of matrix-oriented NADH active site of complex I for activity measurements, purified mitochondria and mitochondrial membranes from cybrid samples were disrupted by osmotic shock through dilution to 2 mg protein/ml in 20 mM Tris-HCl, pH 8.5 followed by one freeze-thaw cycle and then kept on ice for further analysis. Enzymatic activities were measured at 25°C using a SpectraMax ABC Plus plate reader spectrophotometer (Molecular Devices). In general, enzyme activities for each biological sample were determined in three technical replicates.

NADH oxidase (complex I) activity was measured spectrophotometrically for 3 min at 340 nm by adding mitochondrial membranes at final protein concentrations of 70 and 140 μg protein·ml^-1^ for control and ND3^S45P^ mutant cybrid samples, respectively, to the reaction mixture containing 150 μM NADH, 15 µM horse-heart cytochrome *c*, 20 mM Tris-HCl, pH 8.5 and by following the linear decrease of NADH absorbance (*ε*_340_ = 6.22 mM^−1^·cm^−1^). To correct for background activities from other NADH oxidizing enzymes in the sample, inhibitor insensitive rates were determined in the presence of 10 µM complex I inhibitor, *n*-decyl-quinazoline-amine (DQA), and subtracted from total activities. All NADH oxidase activities were corrected for variations in complex I content between samples.

The content of complex I in mitochondrial membranes was assessed by measuring NADH oxidation (ε_340 nm_= 6.22 mM^−1^·cm^−1^) in the presence of the artificial electron acceptor hexaammineruthenium(III)chloride (HAR). This artificial acceptor specifically measures the transhydrogenase activity of complex I-bound flavine mononucleotide and does not depend on a functional ubiquinone reduction site. To start the reaction, 100 µl of mitochondrial membranes (25 μg protein/ml) in 2 mM HAR, 500 µM NaCN, 2 µM antimycin A, 20 mM Tris-HCl, pH 8.5 were mixed with 100 µl of 200 µM NADH.

As a measure of mitochondrial content, citrate synthase (CS) activity was used. CS activity was measured essentially as described by (Haas *et al*, 1995; Shepherd & Garland, 1969; Spinazzi *et al*, 2012) and is based on the formation of 5-thio-2-nitrobenzoate ions followed spectrophotometrically (ε_412 nm_= 13.6 mM^−1^·cm^−1^). The reaction was started by adding 0.5 mM freshly prepared oxaloacetate to mitochondrial membranes (12.5 μg protein·ml^−1^) in 0.05 mM 5,5′-dithiobis-2-nitrobenzoic acid (DTNB), 0.15 mM acetyl coenzyme A, 0.1% Triton X-100, and 100 mM Tris-HCl, pH 8.0.

### Inhibitor titrations

Two different types of specific complex I inhibitors, 0.1 nM-100 nM *n*-decyl-quinazoline-amine (DQA) and 1 nM-1 µM rotenone, were used to compare inhibitor-sensitivities in as prepared mitochondrial membranes of control and mutant ND3^S45P^cybrids. Both inhibitors were dissolved in dimethyl sulfoxide (DMSO) at a concentration that was kept constant at 0.01% (v/v) in every reaction mixture. NADH oxidase activities of mitochondrial membranes of control and ND3^S45P^ mutant (at 70 and 140 µg protein/ml, respectively) were measured in the presence of 187 µM NADH and 14.25 µM cytochrome *c* as described above. All NADH oxidase activity rates for all samples studied were corrected for unspecific residual activities determined at 100 nM DQA. Additionally, for rotenone, all rates were corrected for unspecific residual activities measured at 1 µM and 500 nM rotenone for control and ND3^S45P^ mutant, respectively. IC_50_ values for rotenone and DQA were determined as the effective amount of inhibitor per mg mitochondrial protein resulting in the reduction of the inhibitor-sensitive NADH oxidation rate to 50% of the averaged uninhibited rate.

### Deactive to active transition

Deactivation of mammalian complex I from its active A-form into the reversibly deactivated D-form is a temperature-dependent process occurring in the absence of substrates (Kotlyar & Vinogradov, 1990). The D-form is sensitive to divalent cations and sulfhydryl reagents. To convert the enzyme into the D-form, mitochondrial membranes at 2 mg protein/mlin 20 mM Tris-HCl, pH 8.5 were subjected to one freeze-thaw cycle and then incubated at 37°C for 45 min under constant shaking. This resulted in almost complete deactivation of complex I. NADH oxidase activities were measured at 25°C or, when required, at temperatures increasing in 2-degree intervals from 19-35°C using a PerkinElmer Lambda 25 UV/VIS spectrophotometer. Mitochondrial membranes were applied at 25-50 µg protein/ml for control and 50-100 µg/ml for ND3^S45P^ mutant sample in 20 mM Tris-HCl, pH 8.5. The reaction was started by adding 120 µM NADH and 20 µM cytochrome *c* ensuring that the final volume of the reaction mixture was kept constant for all measurements. Oxidation of NADH (*ε*_340_ = 6.22 mM^−1^·cm^−1^) by complex I was recorded for 5 min. Activation occurs with a lag phase that is more pronounced at lower temperatures. To allow for complex I activation and to follow the underlying reconversion (D to A transition), measurements were performed in the absence of divalent cations. This revealed the NADH oxidase activity of total amount of complex I, i.e. the complex I fraction that was always present in the A-form and that fraction of complex I in the D-form that was reconverted to the A-form with initiation of the NADH oxidation measurement. To be able to determine the fraction of enzyme in the A-form in membranes not subjected to the deactivation protocol (called “as prepared”), enzyme activities were measured in the absence of 5 mM MgCl_2_, but also in its presence to lock the fraction of enzyme already in the D-form. Activities in the presence of divalent cations were used to estimate the portion of the enzyme in the A-form by subtraction from the overall activity in the absence of divalent cations. All activities were normalized to HAR activity.

The Arrhenius equation can used to calculate apparent activation energies from the temperature dependence of reaction rates. This approach has been applied previously to characterize the D-to-A transition of complex I (Maklashina *et al*, 2003). Upon reactivation of fully deactivated complex I, the duration of the lag phase depends on the temperature and can thus be used to determine the rate of re-activation and thereby the apparent activation barrier for the deactive to active transition. The time the enzyme needs to transition to full activity in the assays is used to determine *k,* the rate of conversion from the D- to the A-form per minute for each temperature. By linearizing the Arrhenius equation:

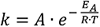

where *A* is the frequency factor, *R* the universal gas constant (8.314 J·K^−1^·mol^−1^) and *T* the temperature in Kelvin, the energy of activation *E_A_* in J·mol^-1^ is obtained from a plot of *ln k* against the reciprocal temperature (1/T) as the negative of the slope times R.

### Oxygen consumption in isolated mitochondria

Oxygen consumption rates (OCRs) were measured at 25°C using a high-resolution respirometry system (Oroboros^TM^ Oxygraph-2k). The oxygen sensor was calibrated with respiration buffer (230 mM sucrose, 5 mM MgCl_2_, 0.5 mM EDTA, 20 mM Tris-HCl, 10 mM phosphate, pH 7.2) at ambient pressure and under constant stirring at 750 rpm at 25°C until a stable signal was obtained (Gnaiger, 2009). Respiration buffer (2 ml) containing control (0.2 mg protein·ml^−1^) or mutant mitochondria (0.3 mg protein·ml^−1^) were placed in the measuring chambers and complex I-linked respiratory states were recorded by making additions in the following sequence: 5 mM pyruvate, 5 mM malate, and 5 mM glutamate (controlled state); 1 mM ADP (active state); 2 μg ml^−1^ oligomycin (ATP synthase-inhibited state); several additions of 50-100 nM carbonyl cyanide *m*-chlorophenyl hydrazone (CCCP) until respiration was maximal (uncoupled state); 1 μM DQA (inhibited state at the level of complex I). To measure complex II-linked respiration, complex I was inhibited by 1 μM DQA prior to the experiments and the respiratory states were determined in the same sequence, except that 10 mM succinate was used to initiate the controlled state and 10 mM malonate was used to obtain the respiratory chain inhibited state.

To determine P/O ratios as a measure of oxidative phosphorylation efficiency (Salin *et al*, 2018), the amount of oxygen needed to phosphorylate 75 nmol ADP ·ml^−1^ in the active state was determined under otherwise identical conditions. Complex I- and complex II-linked respiration were initiated subsequently in the same experiment. Complex I-linked respiration was measured in the presence of complex I substrates and after two consecutive 75 nmol ADP·ml^−1^ additions. To switch the electron input of the respiratory chain, complex I was inhibited by 1 μM DQA and 10 mM succinate was added.

All oxygen consumption rates are expressed as nmol O_2_·min^-1^·mg^-1^ protein and were corrected for mitochondrial content assessed by citrate synthase activity as described above. Respiratory control ratios were calculated by dividing active or uncoupled rates by controlled rates.

### High-resolution respirometry measurements in intact cells

Oxygen consumption rates (OCRs) were measured at 37°C using a high-resolution respirometry system (Oroboros^TM^ Oxygraph-2k). Before each measurement, the oxygen sensor was equilibrated with the growth medium containing glucose (DMEM 4.5 g glucose·l^−1^ medium plus 10% FBS) and internal zero-oxygen calibration was performed. Oxygen concentration and flux were simultaneously recorded and analyzed using the proprietary DatLab software.

Freshly harvested cells were resuspended in 2 ml growth medium mentioned above (final cell concentration: 2 × 10^6^·ml^−1^), placed in the sealed chamber and stirred at 750 rpm. The initial basal rate of respiration without any addition represents the physiologically coupled steady-state fed by endogenous substrates. Subsequently, inhibiting ATP synthase by adding 1 µg·ml^−1^ oligomycin yields a residual non-phosphorylating rate representing the proton leak of the inner mitochondrial membrane. Stepwise addition of 1 μM CCCP results in the uncoupled rate representing maximal uncoupled activity of the respiratory chain. Finally, complex I was inhibited with 5 μM rotenone and mitochondrial respiration was inhibited by 2.5 μM antimycin A. This residual non-mitochondrial respiration was subtracted from all rates. From these experimentally determined rates, respiration-linked ATP production was calculated as difference between basal and leak rates. The spare respiratory capacity reflecting the ability of cells to respond to the energetic demand was obtained as the difference between uncoupled and basal rates. Oxygen consumption rates are expressed as nmol O_2_.min^-1^·10^-6^ cells and normalized to citrate synthase activity that was measured spectrophotometrically with 25 × 10^4^ cells·ml^−1^ using the CS activity assay described above.

### Analysis of mitochondrial morphology

20.000 cybrids/cm^2^ and 5.000 fibroblasts/cm^2^ were seeded on 15.6 mm coverslips (Falcon) and grown in glucose or galactose medium. Cells were fixed with 4% formaldehyde (Sigma) for 20 min and permeabilized with 0.2% Triton X-100 for 10 min. Then samples were blocked with 10% goat serum (GS, Euroclone), incubated with a mouse monoclonal anti-TOM20 primary antibody (1:1000, Santa Cruz, sc-17764) in 0.1% GS for 2 hours, followed by incubation with goat anti-mouse IgG Alexa Fluor 488 secondary antibody (1:1000, ThermoFisher, A11029) and Alexa Fluor 555 Phalloidin (1:50, ThermoFisher, A34055) in 0.1% GS for 1 hr. Nuclei were counterstained using DAPI and mounted using ProLong Gold Antifade Reagent (Cell Signaling). 80-120 cybrids and 68-119 fibroblasts were acquired using the Plan-Apochromat 63X/1.40 Oil DIC M27 objective of an Axio-Imager M2 microscope equipped with Apotome 2 (Carl Zeiss Microscopy). The morphology of mitochondria was quantified using the macro Mitomorph (Yim *et al*, 2019) for ImageJ (National Institutes of Health, Bethesda). A cybrid/fibroblast was classified as with “network” mitochondria or “no network” mitochondria, if mitochondria were classified by Mitomorph as network or not in at least 50%/60% of the total mitochondria of the cell. The remaining cells were classified as having “intermediate” mitochondria.

### Electron microscopy

8.500 cybrids/cm^2^ were plated on Aclair foils (Plano) and after 24 hours were fixed for 1 hour with pre-warmed 2% glutaraldehyde (Sigma), 2.5% sucrose, 3 mM CaCl_2_, and 100 mM Hepes (pH 7.4). Then, sample were washed with with 0,1M cacodylate (Sigma) and postfixed with 1% OsO_4_ (Sigma), 10 mg/ml potassium ferrocyanide, 1.25% sucrose, and 100 mM sodium cacodylate (pH 7.4) for 1 hour on ice. Then, cells were dehydrated using 50%, 70%, 90%, and 100% ethanol, and embedded in Epon resin (Fluka), trimmed and stained with uranylacetate for 15 min. Samples were observed under a transmission electron microscope (JEOL JEM2100PLUS) equipped with a GATAN OneView camera.

### Mass spectrometric analysis of NAD^+^, NADH and CoenzymeA (CoA)

Control and ND3^S45P^ cybrids were grown in triplicates on 10 cm^2^ dishes in presence of glucose or galactose for 48 hours reaching a final concentration of 3 million cells/dish. Then, samples were washed in PBS for 2 times, incubated for 20 min on dry ice with pre-cooled extraction solution (50% LC-MS grade methanol, 30% LC-MS grade acetonitrile, and 20% LC-MS grade water), scraped and immediately processed. NAD^+^, NADH and CoA were quantified by Liquid Chromatography coupled to Electrospray Ionization Tandem Mass Spectrometry (LC-ESI-MS/MS) using previously described procedures (Liu *et al*, 2018; Sasaki *et al*, 2020) with several modifications:

For metabolite extraction, pre-cooled methanol/acetonitrile/water 5:3:2 (v/v/v) was added to the cells in the culture dish (300 µl per 1 × 10^6^ cells), followed by incubation for 20 min on dry ice. The resulting cell suspension was transferred to tubes containing ceramic beads. Cells were homogenized using the Precellys 24 Homogenisator at 6,400 rpm (twice 10 sec with 5 sec break) and directly put on ice again. To 300 µl of homogenate 10 µl of the isotope-labeled internal standard ^13^C_5_-NAD^+^ (100 µM in water, Eurisotop) were added. Metabolites were extracted using a pre-cooled ThermoMixer (Eppendorf) at 4 °C and 900 rpm for 20 min. After centrifugation (16,100 RCF, 20 min, 4 °C), the supernatant was dried under a stream of nitrogen, and the residue was resolved in 100 µl of water. After mixing and centrifugation, 90 µl of supernatant were transferred to autoinjector vials and immediately measured (Liu *et al*, 2018).

10 µl of sample were injected onto an Atlantis T3 column (150 mm × 2.1 mm ID, 3 µm particle size, Waters) and detection using a QTRAP 6500 triple quadrupole/linear ion trap mass spectrometer (SCIEX). The liquid chromatograph (Nexera X2 UHPLC System, Shimadzu) was operated at 40°C and a flow rate of 0.15 ml/min with a mobile phase of 5 mM ammonium formate in water (solvent A) and methanol (solvent B). Metabolites were eluted with the following gradient: initial, 0 % B; 10 min, 70 % B; 15 min, 70 % B; 16 min, 0 % B; and 20 min, 0 % B (Sasaki *et al*, 2020). Metabolites were monitored in the positive ion mode with their specific Multiple Reaction Monitoring (MRM) transitions (**Table S3**). The instrument settings for nebulizer gas (Gas 1), turbogas (Gas 2), curtain gas, and collision gas were 60 V, 40 V, 40 V, and high, respectively. The Turbo V ESI source temperature was 450°C, and the ion-spray voltage was 5.5 kV. The values for declustering potential (DP), entrance potential (EP), collision energy (CE), and cell exit potential (CXP) of the different MRM transitions are listed in **Table S3**.

The quantifier peaks of endogenous NAD^+^, NADH, CoA and the internal standard ^13^C_5_-NAD^+^ were integrated using the MultiQuant 3.0.2 software (SCIEX). The endogenous metabolites were quantified by referring their peak areas to those of the internal standard. The calculated amounts of NAD^+^, NADH and CoA were normalized to the protein content of the sample.

### Complexome profiling

Complexome profiling was performed essentially as described earlier (Heide *et al*, 2012): For separating protein complexes by blue native electrophoresis (Wittig *et al*, 2006) isolated mitochondria were pelleted and resuspended in ice-cold solubilization buffer (50 mM NaCl, 2 mM 6-aminohexanoic acid, 1 mM EDTA, 50 mM imidazole-HCl, pH 7.0). A total of 25 complexome profiling experiments were performed with 2-3 independent replicates of each sample type (**Table S4**). Purified mitochondria were used for cybrids grown on glucose, but since in comparison to existing datasets using crude mitochondria no significant effects on the relative abundance of mitochondrial proteins or improvements in protein detection were observed, the final purification step was omitted for cybrids on galactose and fibroblasts. Samples were solubilized by digitonin (SERVA, 19551) at detergent: protein ratios of 6:1 g/g. Supernatants were recovered by centrifugation at 22,000 × *g* for 20 min at 4°C and supplemented with Coomassie brilliant blue G-250 (SERVA, 17524) loading buffer (5% Coomassie blue, 500 mM aminocaproic acid and 50% glycerol). 200 µg protein per lane was loaded on a 4%–16% polyacrylamide gradient and electrophoresis was started at 100 V for 30 min and continued for 10 hours after exchanging the cathode buffer setting the power supply to 100 V, 30 mA, 10 W at 4–6°C. After electrophoresis, gels were incubated in fixing solution (50% methanol, 10% acetic acid, 100 mM ammonium acetate) for 60 min, followed by staining with 0.025% Coomassie brilliant blue G-250 in 10% acetic acid for 2-3 h. For destaining, gels were incubated 2-3 times for 1 h in 10% acetic acid, followed by two washing steps and storage in deionized water. To prepare a template for cutting, a full-size color image was captured using a flatbed ImageScanner III (GE Healthcare).

In-gel trypsin digestion was performed according to the method outlined in (Emes *et al*, 2008; Heide *et al*, 2012) with minor modifications. Gel lanes were divided into 60 equal slices (∼2 mm), starting at the bottom of the gel. Each gel slice was diced into smaller pieces and then transferred to a 96-well filter microplate (Millipore, MABVN1250) fitted into to a 96-well plate (MaxiSorp^TM^ Nunc) to collect waste solutions. Wells were prefilled with 200 µl of 50% methanol, 50 mM ammonium hydrogen carbonate, pH unadjusted (AHC). To completely remove Coomassie dye, the gel pieces were washed with AHC three times for 30 min at room temperature under gentle agitation until the flow through turned colorless. The remaining solution was removed by centrifugation (1000 x *g*, 15 s) followed by a quick rinse with 200 µl of 50 mM AHC. In the next step, gel pieces were incubated with 120 µl of 10 mM DL-dithiothreitol (Sigma, 43815) in 50 mM AHC for 60 min at room temperature. After a short spin (1000 x *g*, 15 s, RT), 120 µl 30 mM 2-chloroacetamide (Sigma, 22790) in 50 mM AHC was added to each well for cysteine alkylation and incubated for another 45 min in the dark. The solvent was removed by a quick spin and gel pieces were dehydrated by a short incubation (15 min) in 200 µl 50% methanol, 50 mM AHC. Excess solution was discarded and gel pieces were dried at room temperature for 30-45 min followed by an incubation at 4°C for 30 min with 20 µl 5 ng/µl trypsin (sequencing grade Promega, V5111) in 50 mM AHC, 1 mM CaCl_2_ (pH unadjusted). Then, 50 µl 50 mM AHC was added to cover the gel pieces for incubation overnight at 37°C. The following day, the peptide supernatants were collected by centrifugation (1000 x *g*, 1 min) onto a 96-well PCR microplate (Axygen). Finally, gel pieces were washed once more with 50 µl of 30% acetonitrile (ACN), 3% formic acid (FA) for 20 min and the remaining peptides were collected by short spin centrifugation. Finally, the solution containing peptides were dried completely in a SpeedVac Concentrator Plus (Eppendorf) for 3–4 h at 45°C and resuspended in 20 µl of 5% ACN, 0.5% FA. Plates were stored at –20°C for subsequent analysis with LC-MS/MS. All solutions for this procedure were prepared with MS/HPLC grade water and chemicals.

The settings for the mass spectrometer operation were the same as described previously (Guerrero-Castillo *et al*, 2017). Peptides were separated by reverse phase liquid chromatography (LC) and analyzed online by tandem mass spectrometry (MS/MS) in a Q-Exactive Orbitrap mass spectrometer equipped with an Easy nLC1000 nano-flow ultra-high-pressure LC system (Thermo Fisher Scientific) at the front end using 100 μm ID × 15 cm length PicoTip™ EMITTER columns (New Objective, FS360-100-8-N-5-C15) filled with ReproSil-Pur C18-AQ reverse-phase beads (3 μm particle size and 120 Å pore size; Dr. Maisch GmbH, Germany) with 30 min linear gradients of 5–35% acetonitrile, 0.1% formic acid, followed by 35–80% ACN, 0.1% FA (5 min) at a flow rate of 300 nl/min and a final column wash with 80% ACN (5 min) at 600 nl/min. The mass spectrometer was operated in positive ion mode switching automatically between MS and data-dependent MS/MS of the top 20 most abundant precursor ions. Full-scan MS mode (400–1400 *m/z*) was set at a resolution of 70,000 *m*/*Δm* with an automatic gain control target of 1 × 10^6^ ions and a maximum injection time of 20 ms. Selected ions for MS/MS were analyzed using the following parameters: resolution 17,500 *m*/*Δm*, automatic gain control target 1 × 10^5^; maximum injection time 50 ms; precursor isolation window 4.0 Th. Only precursor ions of charge *z* = 2 and *z* = 3 were selected for collision-induced dissociation. The normalized collision energy was set to 30% at a dynamic exclusion window of 60 s. A lock mass ion (*m/z* = 445.12) was used for internal calibration (Olsen *et al*, 2005).

Raw MS data files from all slices of the 25 profiles were analyzed using MaxQuant version 1.5.0.25 (Cox & Mann, 2008) using the settings previously described (Huynen *et al*, 2016). For protein group identification, peptide spectra were searched against reviewed *Homo sapiens* proteome obtained from UniProt database (ID: UP000005640, April 2021) and sequences of pig trypsin and a list of known contaminants e.g. BSA and human keratins. In addition to the original sequence of subunit ND3, an entry was added manually in which serine 45 was changed to proline. Standard parameters for the searches were modified slightly. N-terminal acetylation and oxidation of methionine were set as dynamic modifications and cysteine carbamidomethylation as the fixed modification. Trypsin was selected as protease and up to two missed cleavages were permitted for trypsin digestion. Matching between runs was allowed and matching time window was set at 2 min. False discovery rate (FDR) as determined by target-decoy approach, the limit was strictly set to 0.01. Minimal peptide length was fixed at six residues. Abundance comparisons between samples were achieved by applying label-free quantification using the obtained intensity-Based Absolute Quantification (iBAQ) values. Variations driven from MS sensitivity and protein quantity among runs were corrected by normalizing against the sum of total iBAQ values from each sample. Data were normalized for the sum of iBAQ values of mitochondrial proteins annotated in MitoCarta 3.0 from each sample, in total 1047 mitochondrial proteins were identified in the samples. The migration patterns of the identified proteins were hierarchically clustered by an average linkage algorithm with centered Pearson correlation distance measures using the Cluster 3.0 algorithm (de Hoon *et al*, 2004). The resulting complexome profiles consist of a list of proteins clustered according to the similarity of their migration patterns in blue native electrophoresis and were illustrated as heatmaps representing the normalized abundance in each gel slice by a three-color gradient (black/yellow/red). The molecular masses were calibrated for each profile using the known apparent molecular masses for membrane-bound and water-soluble protein complexes (**Figure S4**;(Wittig *et al*, 2010). Further manual clustering and analysis were carried out in Microsoft Excel.

The sum of the abundances in all slices of the migration profiles was used to calculate the relative changes in the overall abundances of each protein. Abundance changes in multiprotein complexes were calculated by averaging the profiles of all subunits reliably detected by mass spectrometry unless otherwise specified. Values were renormalized again against mutation impact and carbon source.

### Software and statistical analysis

Versions and sources of the software packages used are listed in **Table S5**. Statistical analysis was performed using the GraphPad Prism 8.4.3 software package (GraphPad Software, Inc., San Diego, California, USA). Data were evaluated using unpaired two-tailed Student’s *t*-tests or two-way ANOVA with Sidak correction to assess the abundance of multiprotein complexes. Curves were fitted by nonlinear or linear regression without weighting. Values are given as means ± S.D. with *n* being the number of independent biological samples. Significance is indicated as follows: \**p* < 0.05, \*\**p* < 0.01, \*\*\**p* < 0.001, \*\*\*\**p* < 0.0001.

## Data availability

The complexome profiling datasets were deposited in the CEDAR database, accession code CRXxx (www3.cmbi.umcn.nl/cedar/browse/experiments/CRX32; (van Strien *et al*, 2021)).

## Acknowledgements

This work was supported by TOP project 714.017.004 of the Netherlands Organization for Scientific Research (NWO), by TOP project 91217009 of the Netherlands Organization for Health Research and Development, and Collaborative Research Center 1218 (Project 269925409) of the German Research Foundation (DFG). We would like to thank Coby Laarakkers and Erica Barry for their support with some of the experiments.

## Author contributions

U.B. and S.A. designed the study and interpreted the results; Z.A.A. performed most of the experiments, developed functional assays and interpreted the results; A.C.O, helped with development and optimization of functional assay and contributed to the complexome profiling analyses; M.Z., E.B., E.I.R. and S.B. performed microscopic and metabolomic analyses; R.J.R. provided fibroblasts and muscle biopsies; U.B., Z.A.A. and S.A. wrote the paper and all authors read, corrected and approved the final version.

