## Supplementary Material for "Pathogenesis of mtDNA point mutation m.10191T>C affecting complex I function is a multifactorial process leading to metabolic remodeling of mitochondria"

for

**Figures S1-S4**

**Tables S2-S5**

**Supplementary Table S1 is provided as MS-Excel file.**

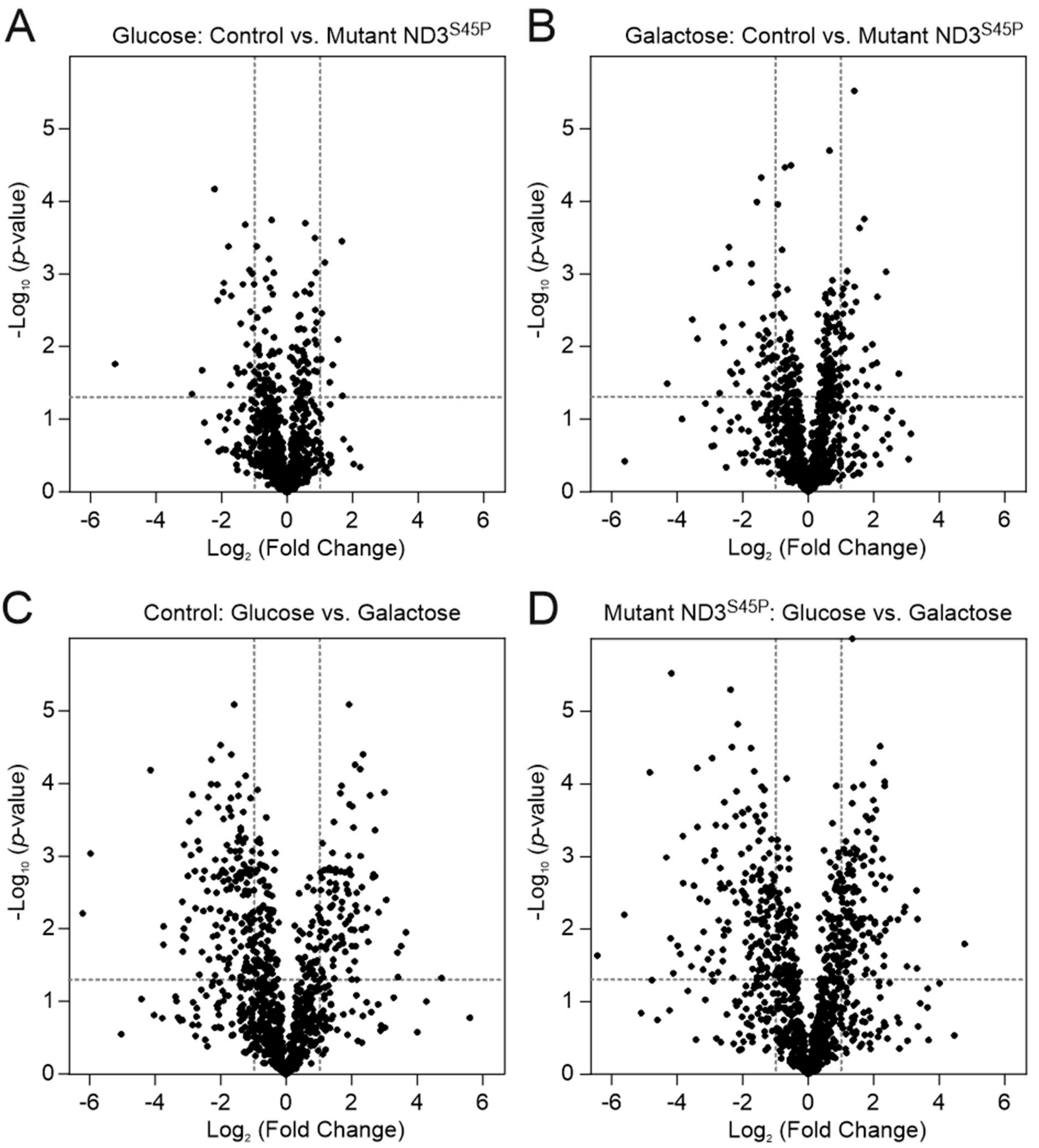


**Figure S1:** *Remodeling of the mitochondrial proteome.* Volcano plots illustrate the fold change in abundance of ~1000 mitochondrial proteins induced by mutation $ND3^{S45P}$ in cybrids on glucose or galactose media. Total abundance of each protein was calculated by label-free quantification in complexome profiling datasets from three independent batches of mitochondria each. Impact of mutation $ND3^{S45P}$ on glucose (**A**) and galactose (**B**) media; impact of switching control (**C**) and $ND3^{S45P}$ (**D**) cells from glucose to galactose media.

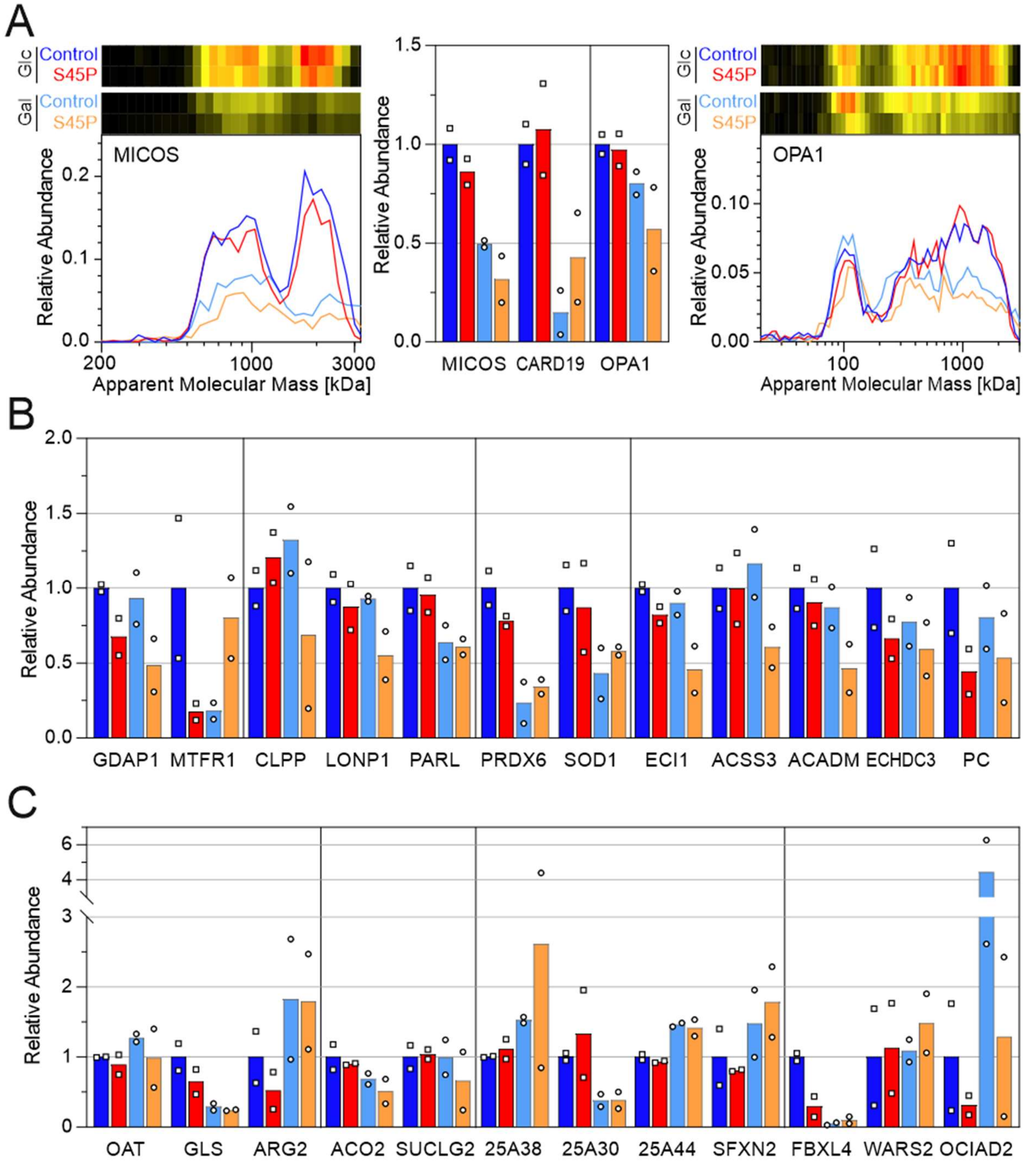


**Figure S2:** *Impact of mutation ND3$^{S45P}$ and forced OXPHOS on the migration profiles and abundances of selected mitochondrial proteins in patient (S45P) and control fibroblasts.* **A**, migration profiles with corresponding heatmaps and relative abundances of MICOS (average of MIC60, MIC13, MIC19 and MIC26), CARD19 and OPA1; **B**, abundances of selected proteins involved in mitochondrial dynamics (GDAP1, MTFR1), quality control (CLPP, LONP1, PARL), oxidative defense (PRDX6, SOD1) and β-oxidation (ECI1, ACSS3, ACADM, ECHDC3, PC); **C**, selected amino acid metabolism enzymes (OAT, GLS, ARG2), TCA cycle components (ACO2, SUCLG2), mitochondrial transporters (SLC25A38, SLC25A30, SLC25A44, SFXN2) and disease related proteins (FBXL4, WARS2, OCIAD2). Relative changes for the indicated proteins are shown as bar graphs. Color code: control cells (C1, C2), dark/light blue on glucose/galactose media; patient fibroblasts (P1, P2), red/orange on glucose/galactose media. Relative abundances were renormalized for each protein by setting the average of controls on glucose to unity and are presented as mean of two independent control and patient samples.

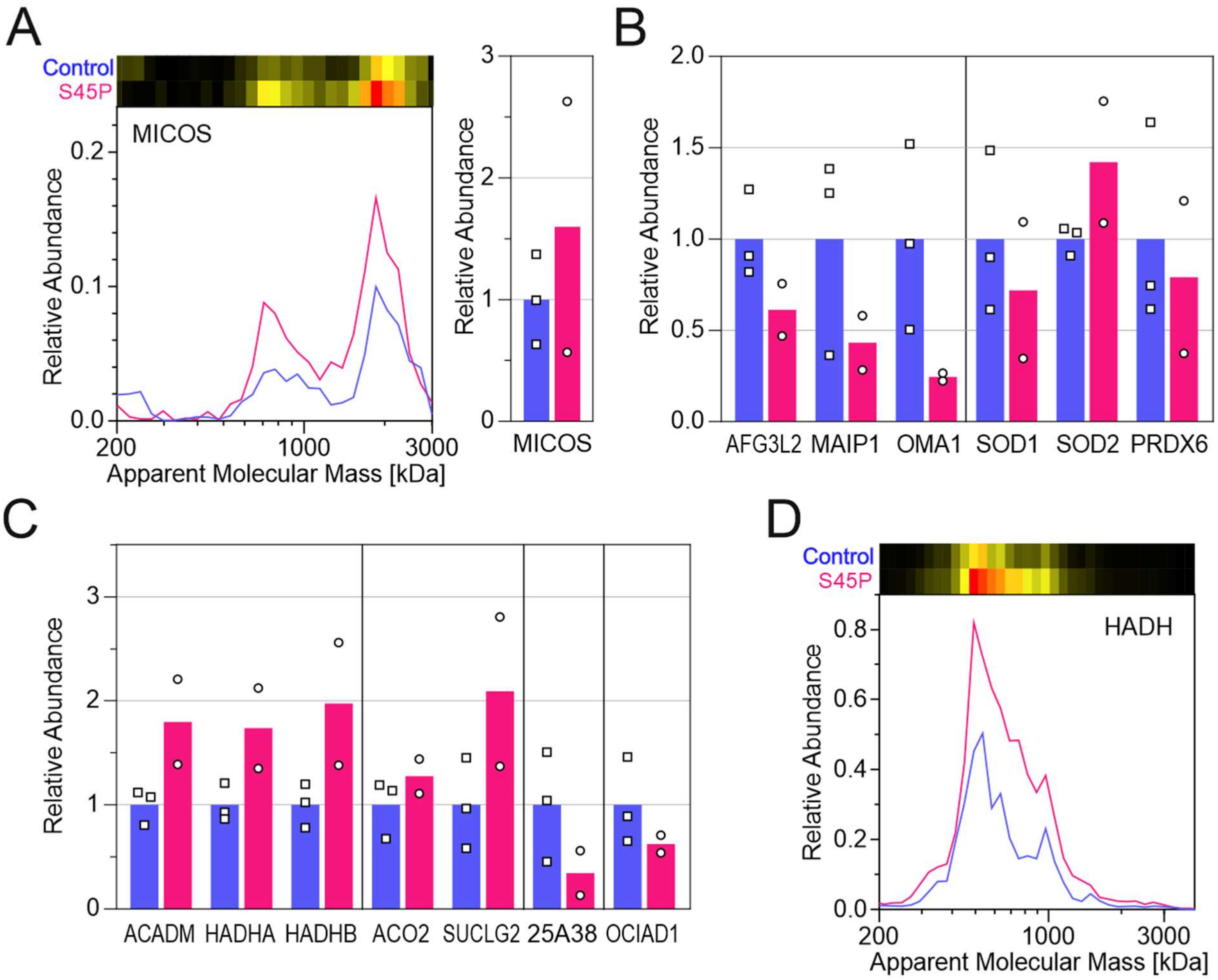


**Figure S3:** *Impact of mutation ND3$^{S45P}$ and forced OXPHOS on the migration profiles and abundances of selected mitochondrial proteins in patient and control muscle biopsies.* **A**, migration profiles with corresponding heatmaps and overall relative abundance of MICOS (average of MIC60, MIC19 and MIC26); **B**, abundances of selected proteins involved in mitochondrial quality control (AFG3L2, MAIP1, OMA1) and oxidative defense (SOD1, SOD2, PRDX6); **C**, abundances of selected proteins involved in β-oxidation (ACADM, HADHA, HADHB), TCA cycle components (ACO2, SUCLG2), mitochondrial transporter SLC25A38 and disease-related protein OCIAD1; **D,** migration profiles with corresponding heatmaps of trifunctional enzyme HADH (average of HADHA and HADHB). Relative changes for the indicated proteins are shown as bar graphs (**A-C**). Color code: control muscle tissues, blue; patient muscle tissues, magenta. Relative abundances were renormalized for each protein by setting the average of controls to unity and are presented as mean of three independent control and two patient samples.

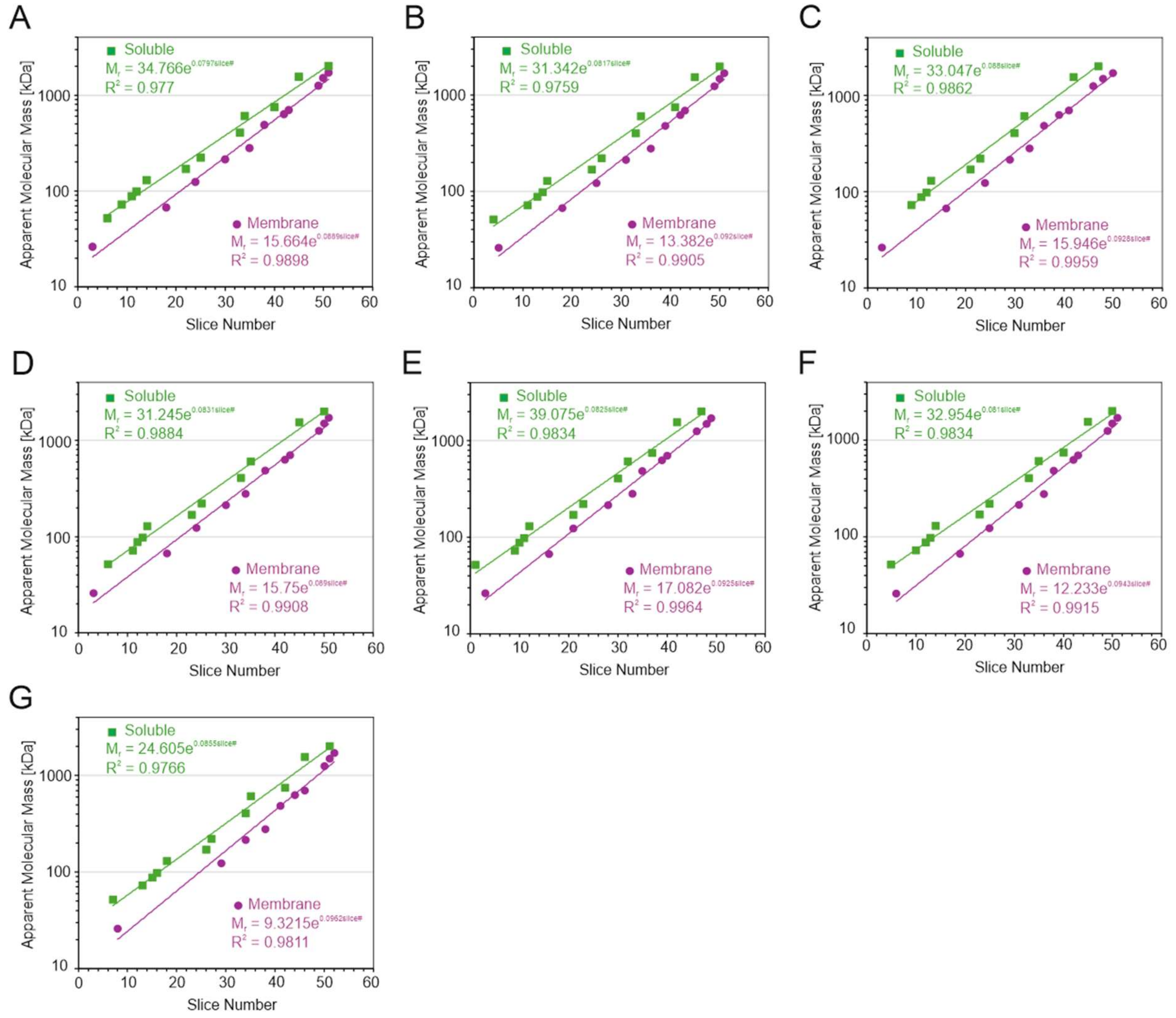


**Figure S4:** *Mass calibration of averaged migration profiles for complexome profiling*. **A**, mass calibration of cybrids grown on glucose for replicate 1 of control and ND3$^{S45}$; **B**, cybrids grown on glucose replicates 2, 3 of control and ND3$^{S45P}$; **C**, cybrids grown on galactose for replicates 1-3 of control and ND3$^{S45P}$; **D**, fibroblasts grown on glucose for C1, C2 and P1, P2; **E**, fibroblasts grown on galactose for C1, P1; **F**, fibroblasts grown on galactose for C2, P2; **G**, muscle biopsies all five samples. The apparent masses for membrane integral (magenta) and water-soluble (green) proteins were separately determined by fitting the following calibration proteins and protein complexes to an exponential curve. Membrane protein standards: MIC27 (26 kDa), TOMM70 (67 kDa), respiratory complex II (123 kDa), complex IV (214 kDa), voltage-dependent anion channel VDACs (279 kDa), complex $III_2$ (485 kDa), complex V monomer (626 kDa), supercomplexes $III_2$-$IV_1$ (699 kDa), complex V dimer (1252 kDa), supercomplexes $S_0$ (1494 kDa) and $S_1$ (1708 kDa). Soluble protein standards: monomeric ATP synthase subunit beta ATP5B (51.7 kDa), complex I assembly intermediates NDUFV1/NDUFV2 (72 kDa), NDUFS1/NDUFA2 (88 kDa), citrate synthase CS (98 kDa), Q module (129 kDa) and Q module with assembly factors NDUFAF3/NDUFAF4 (170 kDa), the homo-tetramer of aldehyde dehydrogenase X (221 kDa), heptameric 60 kDa heat-shock protein HSPD1 (406 kDa), $\alpha_2\beta\gamma$ $NAD^+$ dependent isocitrate dehydrogenase (608 kDa), 20S proteasome (750 kDa), propionyl-CoA carboxylase (1548 kDa), 26S proteasome (2,000 kDa).

**Table S2**: *List of samples from m.10191T>C (p.S45P$^{ND3}$) patients and controls.*

| Individual | Code | Age | Gender | Muscle | Fibroblasts |
|---|---|---|---|---|---|
| | | *Years* | | *% Heteroplasmy* | |
| Control 1 | C1 | 2 | Male | 0 | 0 |
| Control 2 | C2 | 26 | Male | 0 | 0 |
| Control 3 | C3 | 25 | Male | 0 | - |
| Patient 1 | P1 | 2 | Male | 75 | 75 |
| Patient 2 | P2 | 25 | Male | 88 | 82 |

**Table S3**: *MRM transitions and compound-specific parameters*

| Q1 Mass | Q3 Mass | ID | DP | EP | CE | CXP | Quantifier |
|---|---|---|---|---|---|---|---|
| *Da* | | | *V* | | | | |
| 664.1 | 136.1 | $NAD^+$ | 110 | 10 | 35 | 10 | |
| 664.1 | 428.1 | $NAD^+$ | 110 | 10 | 36 | 10 | X |
| 664.1 | 524.1 | $NAD^+$ | 110 | 10 | 27 | 10 | |
| 664.1 | 542.2 | $NAD^+$ | 110 | 10 | 39 | 10 | |
| 666.1 | 302.0 | NADH | 95 | 10 | 49 | 10 | |
| 666.1 | 514.1 | NADH | 95 | 10 | 37 | 10 | X |
| 666.1 | 649.1 | NADH | 95 | 10 | 40 | 10 | |
| 768.1 | 136.1 | CoA | 100 | 10 | 35 | 10 | |
| 768.1 | 159.1 | CoA | 100 | 10 | 35 | 10 | |
| 768.1 | 261.1 | CoA | 100 | 10 | 35 | 10 | X |
| 669.1 | 428.1 | $^{13}C_5$-$NAD^+$ | 110 | 10 | 36 | 10 | X |
| 669.1 | 529.1 | $^{13}C_5$-$NAD^+$ | 110 | 10 | 27 | 10 | |

**Table S4:** *Complexome profiling experiments*

| Sample | Control | Mutant ND3$^{S45P}$ |
|---|---|---|
| Cybrids on glucose | n=3 | n=3 |
| Cybrids on galactose | n=3 | n=3 |
| Fibroblasts on glucose | n=2 | n=2 |
| Fibroblasts on galactose | n=2 | n=2 |
| Muscle biopsy | n=3 | n=2 |

**Table S5**: *Software*

| | |
|---|---|
| MaxQuant 1.5.0.25 | https://maxquant.org/ |
| GraphPad Prism 8.4.3 | https://www.graphpad.com/ |
| SoftMax Pro 7.1 | https://www.moleculardevices.com/ |
| Origin | https://www.originlab.com/ |
| PyMol Molecular Graphics System Version 3.0 | https://pymol.org/ |
